# Constitutive and conditional epitope-tagging of endogenous G protein coupled receptors in *Drosophila*

**DOI:** 10.1101/2023.12.27.573472

**Authors:** Shivan L. Bonanno, Piero Sanfilippo, Aditya Eamani, Maureen M. Sampson, Kandagedon Binu, Kenneth Li, Giselle D. Burns, Marylyn E. Makar, S. Lawrence Zipursky, David E. Krantz

## Abstract

To visualize the cellular and subcellular localization of neuromodulatory G-protein coupled receptors (GPCRs) in *Drosophila*, we implement a molecular strategy recently used to add epitope tags to ionotropic receptors at their endogenous loci. Leveraging evolutionary conservation to identify sites more likely to permit insertion of a tag, we generated constitutive and conditional tagged alleles for *Drosophila* 5-HT1A, 5-HT2A, 5-HT2B, Octβ1R, Octβ2R, two isoforms of OAMB, and mGluR. The conditional alleles allow for the restricted expression of tagged receptor in specific cell types, an option not available for any previous reagents to label these proteins. We show that 5-HT1A and 5-HT2B localize to the mushroom bodies and central complex respectively, as predicted by their roles in sleep. By contrast, the unexpected enrichment of Octβ1R in the central complex and of 5-HT1A and 5-HT2A to nerve terminals in lobular columnar cells in the visual system suggest new hypotheses about their function at these sites. Using an additional tagged allele of the serotonin transporter, a marker of serotonergic tracts, we demonstrate diverse spatial relationships between postsynaptic 5-HT receptors and presynaptic 5-HT neurons, consistent with the importance of both synaptic and volume transmission. Finally, we use the conditional allele of 5-HT1A to show that it localizes to distinct sites within the mushroom bodies as both a postsynaptic receptor in Kenyon cells and a presynaptic autoreceptor.

**Significance Statement:** In *Drosophila*, despite remarkable advances in both connectomic and genomic studies, antibodies to many aminergic GPCRs are not available. We have overcome this obstacle using evolutionary conservation to identify loci in GPCRs amenable to epitope-tagging, and CRISPR/Cas9 genome editing to generated eight novel lines. This method also may be applied to other GPCRs and allows cell-specific expression of the tagged locus. We have used the tagged alleles we generated to address several questions that remain poorly understood. These include the relationship between pre- and post-synaptic sites that express the same receptor, and the use of relatively distant targets by pre-synaptic release sites that may employ volume transmission as well as standard synaptic signaling.

## Introduction

Serotonin (5-HT) regulates a variety of behaviors in *Drosophila*, and while flies do not synthesize noradrenaline, the structurally related molecule octopamine (OA) serves many of the same roles (Maqueira et al., 2005; Roeder, 2005). Identifying the cellular and subcellular expression patterns for 5-HT and OA receptors is essential to understand their function and to complement existing connectomic and transcriptomic studies, however the lack of high-affinity antibodies has hampered such investigations. Moreover, antibodies to some other *Drosophila* GPCRs such as mGluR that were previously created (Panneels et al., 2003; Bogdanik, 2004) are now unavailable.

The lack of high-affinity antibodies has made it difficult to determine the distance between presynaptic monoaminergic release sites and their postsynaptic targets in *Drosophila.* In mammals, most if not all aminergic neurotransmitters can signal via “volume” or extra-synaptic transmission, wherein transmitters are released through non-canonical pathways and can diffuse at distances of several microns rather than the nanometer scale of a standard synapse (Agnati et al., 1992; Rice, 2000; Fuxe et al., 2012; Del-Bel and De-Miguel, 2018; Wildenberg et al., 2021). Importantly, extra-synaptic signaling sites will be missed by scoring canonical synaptic structures in connectomic studies or by molecular methods such as GRASP and trans-Tango that rely on physical proximity between cells (Feinberg et al., 2008; Macpherson et al., 2015; Shearin et al., 2018; Sorkaç et al., 2023). Thus, in the absence of high-affinity antibodies, extra-synaptic transmission remains poorly characterized in *Drosophila* despite extensive efforts to map synaptic connectivity in both the larval and adult fly brain.

Another consideration in mapping aminergic pathways is that the same GPCRs can be expressed on closely apposed pre-and postsynaptic neurons (Garcia-Garcia et al., 2014; You et al., 2016; Albert and Vahid-Ansari, 2019), making it difficult to differentiate signal using antibodies to endogenous proteins and light microscopy. Controllable expression of epitope-tagged receptors provides a workaround to this problem.

The use of standardized epitopes also serves as a more general substitute for custom antibodies. Toward this end, several *Drosophila* 5-HT and dopamine (DA) receptors as well as other neuronal proteins have been tagged with GFP at their N- and C-termini (Qian et al., 2017; Alekseyenko et al., 2019; Fendl et al., 2020; Kondo et al., 2020; Certel et al., 2022a, 2022b; Parisi et al., 2023). However, classical studies in mammals indicate that N- and C-terminal tags can disrupt GPCR trafficking (Guan et al., 1992; Dunham and Hall, 2009; Cho et al., 2012), which led us to explore alternative sites. Additionally, although a conditional tag has been reported for one *Drosophila* GPCR (GABA-B-R1) (Fendl et al., 2020), no such reagents have been previously reported for any of the receptors in this study.

We leveraged evolutionary divergence to identify sites in receptor primary amino acid sequences that are more likely to permit insertion of an epitope tag (Sanfilippo et al., 2023), and generated tagged alleles for eight *Drosophila* GPCRs (5-HT1A, 5-HT2B, 5-HT2A, mGluR, Octβ1R, Octβ2R, and two isoforms of OAMB). The expression patterns we observed match the results of previous behavioral studies where available, and generate new hypotheses where data is lacking. Paired with cell-type specific drivers, we used conditional receptor alleles to unambiguously determine their expression in specific neuronal types. We also developed a new marker for serotonergic neurons and observed a surprisingly wide range in its distance from postsynaptic 5-HT receptors across different brain regions. Finally, we used the tagged receptor alleles to demonstrate enrichment in axonal versus dendritic compartments and to differentiate pre-versus postsynaptic 5-HT receptors in the mushroom body.

## Results

### Development of constitutive and conditionally epitope-tagged GPCR alleles

We initially created tagged alleles for six GPCRs (5-HT1A, 5-HT1B, 5-HT2A, 5-HT7, CrzR, and OAMB) where the insertion was placed at the N-terminus, similar to constructs used in the past for mammalian GPCRs (Guan et al., 1992; Andersson et al., 2003; McDonald et al., 2007). We failed to detect expression in most of them, with possible cell-body-restricted expression in two (5-HT1A, 5-HT2B, data not shown). Following studies in mammalian systems showing that preceding an N-terminal tag with a signal sequence partially rescued trafficking defects (Guan et al., 1992; Andersson et al., 2003; McDonald et al., 2007), we then created an additional ten alleles where the tagging cassette was placed immediately after the putative signal sequence, but still close to the N-terminus (mGluR, OAMB, Octβ1R, Octβ2R, Octβ3R, DopEcR, Dop2R, GABA-B-R1, GABA-B-R2, and GABA-B-R3). However, immunolabeling of these alleles was also unsuccessful, despite differences in receptor subclasses and verification of the intended genomic transformations (data not shown). Finally, we used evolutionary conservation of receptor primary amino acid sequences to guide our identification of locations that might be more tolerant of epitope insertions as described previously for ionotropic receptors (Sanfilippo et al., 2023). We analyzed multiple protein alignments for GPCRs between *Drosophila melanogaster* and other *Drosophila* species as well as more distantly related insects. We found that for Class A GPCRs such as 5-HT1A, the 3^rd^ intracellular loop (ICL) is large and often contains amino acid insertions or short repeats that are not conserved. Reasoning that positions that reside within expansion-tolerant stretches, such as G631 in 5-HT1A (Fig. 1A), may also be more tolerant to insertion of an exogenous DNA cassette, we selected similar regions for genetic modification for an additional 6 Class A GPCRs.

**Fig. 1:**
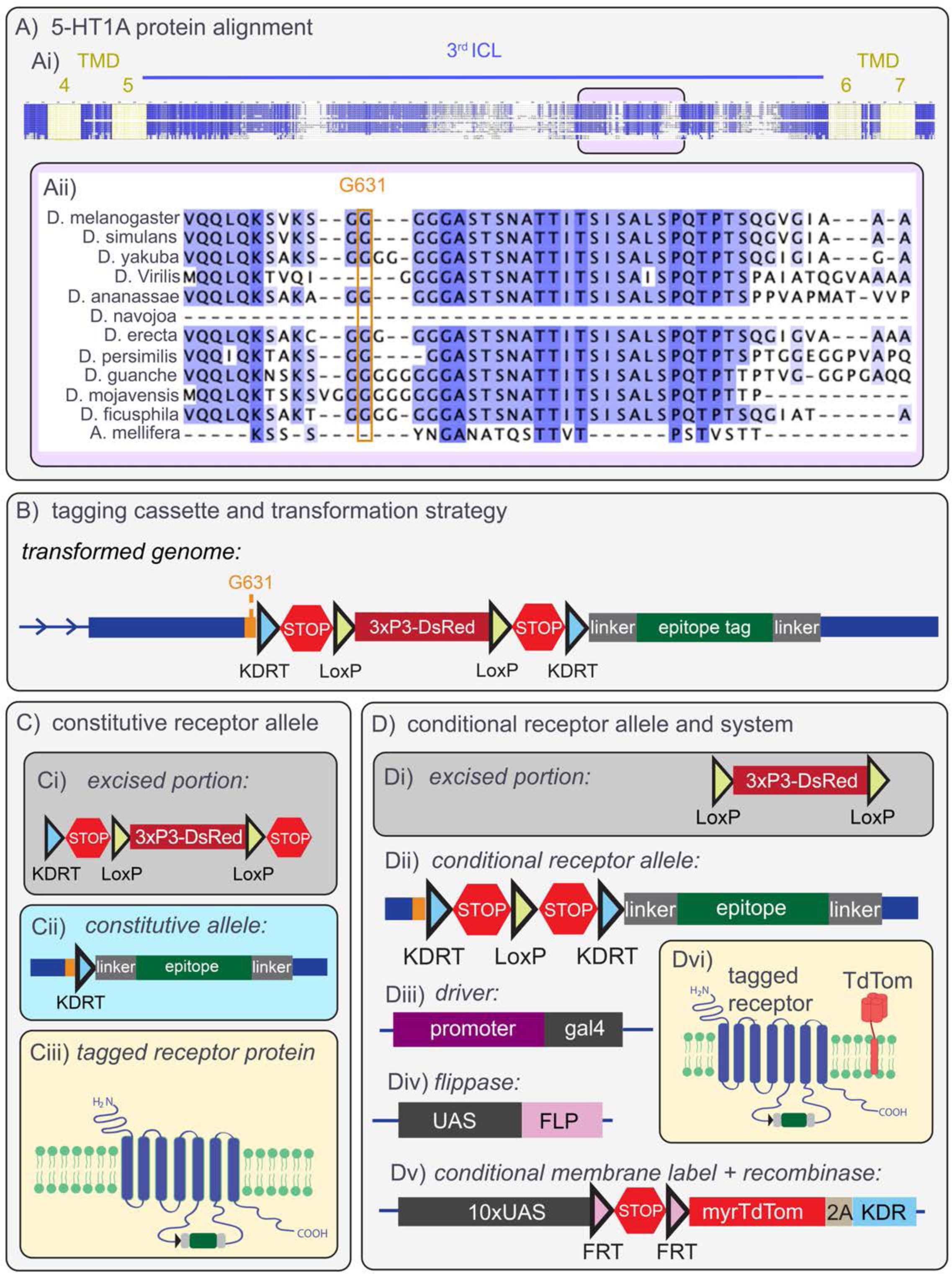
Identification of sites in receptor sequences for epitope tagging and derivation of constitutively and conditionally-tagged alleles. Ai) Selected regions of a multiprotein alignment using 5-HT1A sequences from multiple *Drosophila* species and *Apis mellifera*. Conserved sites are colored blue. Aii) An expanded view of the boxed region in (Ai). The 3^rd^ intracellular loops (ICL) of class A GPCRs, such as 5-HT1A, are large and contain multiple regions of low conservation, as indicated by the absence of blue highlighted residues (e.g., glycine (G) residues downstream of G631). This glycine expansion suggests the region may also tolerate insertion of exogenous protein sequences. B) The DNA insertion for tagging alleles. At baseline, STOP cassettes (or interruption moieties, see text) terminate transcription and translation. The floxed 3xP3-DsRed cassette is used to screen transformants, then removed via recombinases as indicated in panels (C) and (D). Separate KDRT (blue triangles) and LoxP (yellow triangles) recombination sites flank the STOP and 3xP3-DsRed cassettes, respectively. C) Constitutively-tagged receptor alleles are derived from the transformed genome in (B) by crossing to flies expressing KD recombinase (KDR) in the germline. KDR acts on the KDRT sites to excise both the STOPs and the 3xP3-DsRed cassettes (Ci), and yields an in-frame receptor sequence with the epitope insertion (green) flanked by linker sequences (grey) and a residual KDRT site (Cii). Translation of the constitutive allele (Ciii) generates a receptor tagged within the 3^rd^ ICL of the protein. D) Conditional alleles are derived from the tagging construct in (B) by crossing to flies that express Cre recombinase to remove the 3xP3-DsRed cassette (Di), leaving the conditional receptor allele with two STOP cassettes flanked by KDRT sites (Dii). For conditional labeling experiments, the conditional receptor allele (Dii) is crossed to flies expressing a GAL4 driver (Diii), UAS-flippase (Div) and an additional UAS transgene (Dv) that contains both a membrane bound myristoylated fluorophore (myr::TdTom or myr::GFP) and the KDR recombinase, separated by a T2A translational skip element. GAL4 driven by a cell-specific promoter binds to the UAS sites on both of the other (Div and Dv) components of the system. FLP recombinase is thus expressed in the GAL4-defined population and acts on the FRT sites (pink triangles) in (Dv) to remove the STOP cassette. This results in GAL4-dependent expression of both myr::TdT (Dvi, “TdTom”) and KDR. KDR then acts on the KDRT sites (blue triangles) in the conditional receptor allele to remove the STOP cassettes and allow expression of the tagged receptor, resulting in a GAL4-defined population of cells that expresses both the tagged receptor whose spatiotemporal expression pattern is governed by its exogenous genomic locus, and an overexpressed myristoylated fluorophore.

While evolutionary conservation analysis can be applied to any receptor, ideal insertion sites are not always immediately apparent. Class C GPCRs such as mGluR lack a large 3^rd^ intracellular loop and exhibit weak, if any, evidence of evolutionarily tolerated insertions in other regions. Though it is unfortunately no longer available, a monoclonal antibody against *Drosophila* mGluR was previously developed and the epitope binding site was subsequently mapped (Panneels et al., 2003; Bogdanik, 2004). We reasoned that successful labeling using this antibody in the past suggests that its binding site is topographically accessible, and could thus represent a good location for epitope tagging of mGluR and potentially other class C GPCRs.

The receptor tagging cassettes in this study contain either 1xALFA tag (Götzke et al., 2019), or spaghetti-monster V5 (smV5) (Viswanathan et al., 2015) preceded by recombinase-flanked interruption moieties. For each receptor, we derive a constitutively tagged allele in which all regulatory moieties are removed and the tag is in the open reading frame (Fig. 1C), as well as a conditional allele in which interruption moieties/”stop signals” are retained. For the conditional alleles, inclusion of the tag is dependent on orthogonal expression of a recombinase to remove the stop signals (Fig. 1D).

We selected ALFA and V5 epitopes for the receptor alleles generated in this study because commercially available antibodies are of high quality and work well in *Drosophila* tissue. Of these, only ALFA is a recently developed epitope, and we compared the quality of several anti-ALFA antibodies in our immunohistochemical analyses (data not shown). Though the data presented in this study used a mouse Fc-conjugated ALFA nanobody, a similar guinea pig Fc-conjugated nanobody that is available performs equally well and new antibodies should be tested systematically for signal-to-noise ratio as they are made available.

Using ALFA and V5 tags, we generated the following alleles: 5-HT1A-ALFA, 5-HT2B- ALFA, 5-HT2A-smV5, mGluR-ALFA, Octβ1R-smV5, Octβ2R-smV5, OAMB-K3-ALFA, and OAMB-RS-ALFA. None of these localized exclusively to the cell body and all labeled distal neuropils, strongly indicating that trafficking was not impaired as it had been in our initial attempts. We therefore proceeded with more detailed imaging studies of the aminergic class A GPCRs and the single class C representative, mGluR, that we generated.

### Octopamine receptors in the mushroom bodies and central complex

OA pathways are intimately associated with the mushroom bodies (MBs) (Busch et al., 2009; Wu et al., 2013), which are the major locus for learning and memory in insects (Akalal et al., 2006; Modi et al., 2020). Mutation or knockdown of OAMB (Kim et al., 2013) and, more recently Octβ1R (Sabandal et al., 2020), have been proposed to impair learning while knockdown of Octβ2R does not (Sabandal et al., 2020). The expression of OAMB in the MBs is well established by RNA in situ hybridization (Han et al., 1998), custom antibodies (Han et al., 1998; Kim et al., 2013), and GAL4 transgenes (McKinney et al., 2020).

While most *Drosophila* GPCRs have many annotated isoforms, few have been validated by protein or RNA-expression studies. One notable exception is OAMB, which expresses two major protein isoforms (OAMB-K3 and OAMB-AS), both of which have been validated by custom antibodies (Kim et al., 2013). To determine whether alternative isoforms of OAMB could be specifically labeled using our technique, we generated tagged alleles for each splice variant in which the tagging cassette was inserted after the exon specific to each isoform. We find that the tagged versions of both isoforms are expressed in the MB lobes and we detect additional labeling in the EB for OAMB-K3 but not AS. The expression patterns of OAMB-K3-ALFA and OAMB-AS-ALFA (Fig. 2A-B) precisely mimic those reported for the endogenous proteins (Kim et al., 2013). Our success tagging both OAMB variants suggest that a similar strategy might be used to explore the expression patterns of other GPCR splice variants that have been predicted by cDNA analysis but not yet validated by expression studies.

**Fig. 2:**
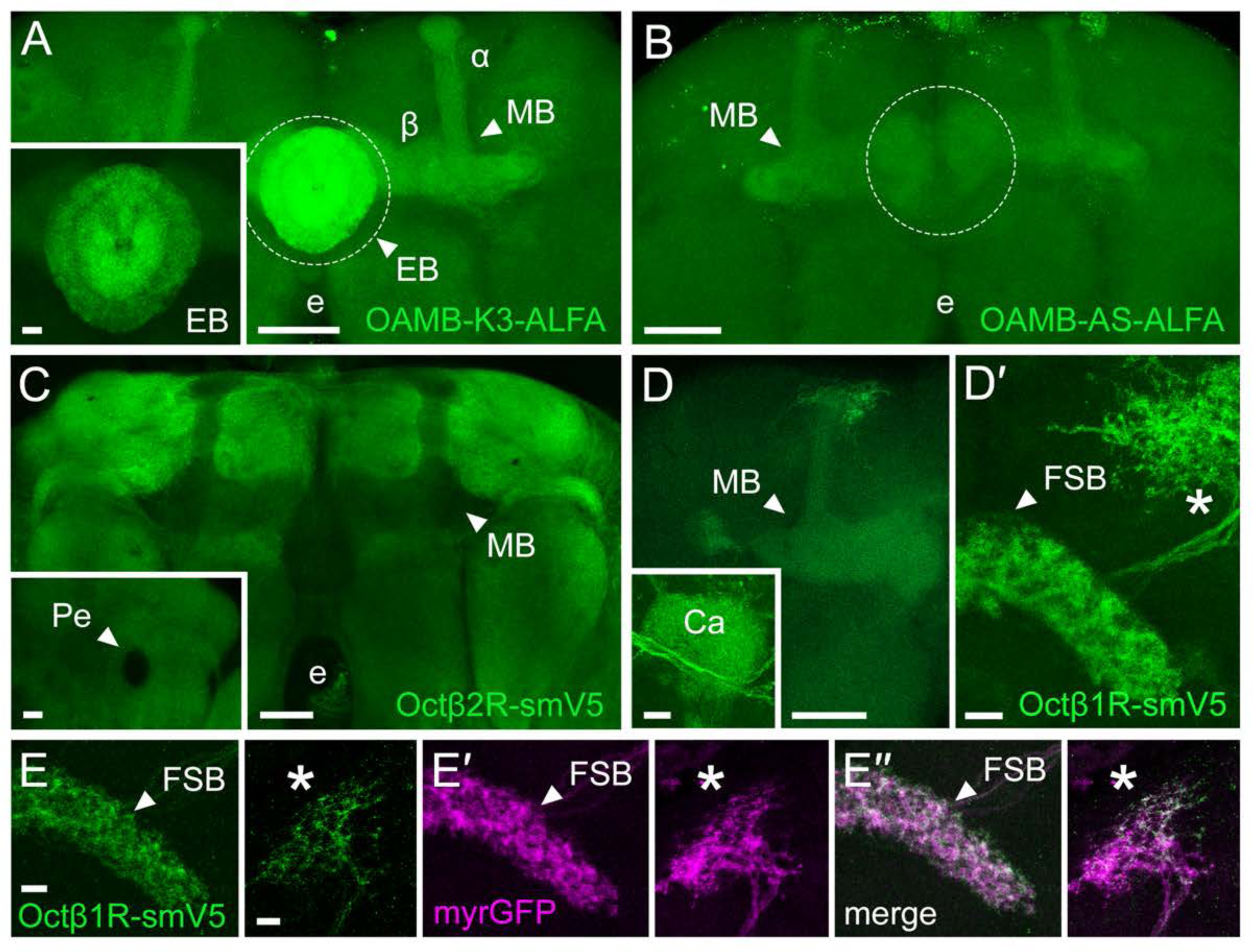
Octopamine receptors OAMB and Oct 1R but not Oct 2R are enriched in the mushroom bodies and central complex. A) OAMB-K3 is enriched in the α/ lobes of the mushroom bodies (MBs), with even more striking enrichment in the ellipsoid body (EB) (inset) that becomes overexposed (dotted circle) when calibrated for the MBs. e: esophagus. B) OAMB-AS is similarly enriched in the α/ lobes of the MBs, but strikingly absent from the EB. C) Oct 2R is expressed throughout the central brain and de-enriched in the MB lobes and peduncle (Pe, inset). D-D′) Oct 1R is expressed in the α/ lobes (D), and the calyx (inset) of the MBs and a dorsal arborization (D′, asterisk), that extends into the fan-shaped body (FSB). E-E′′) The Oct 1R allele (green) conditionally expressed in the 23E10-GAL4 cells localizes to both the dorsal neuropil (asterisk) and the FSB. The conditional expression system simultaneously labels the membrane of these cells with TdTomato (magenta). Scale bars in the main panels of A-D and D′: 50μm; scale bars in A-D and D′ insets and in E-E′′:10μm.

A custom antibody against the C-terminus of Octβ2R was previously reported (Wu et al., 2013) and low levels of expression were observed in the α′/β′ lobes of the MB. Our Octβ2R tagged allele, however, is notably absent from the lobes of the mushroom bodies but broadly expressed throughout the central brain (Fig. 2C).

While GAL4 lines for Octβ1R indicate that it is expressed in the MB (McKinney et al., 2020), reports using custom antibodies or tagged alleles are lacking, and the subcellular localization of the receptor has not yet been investigated. We find that the Octβ1R tagged allele is expressed in both the lobes and calyx of the MB (Fig 2D). Localization at both sites suggest it could mediate OA signaling in the dendrites (the MB calyx) and the axons (the MB lobes) of the KCs.

Octβ1R is also strikingly enriched in a small cluster of neurons that innervates the fan shaped body (FSB) and has an additional dorsolateral projection (Fig. 2D′). Based on the morphology of these projections, we hypothesized that they may represent 23E10 neurons, which have been previously shown to be required for sleep (Donlea et al., 2014, 2018; Pimentel et al., 2016). To test this hypothesis, we used the 23E10 driver to express the conditional Octβ1R allele. We observe labeling identical to that seen with the constitutive allele, demonstrating that Octβ1R is indeed expressed in 23E10(+) neurons (Fig. 2E-E′′). To our knowledge, there is no published RNA-seq for 23E10 cells. Thus, our data constitutes the first report of expression of an OA receptor in this population of cells, and raise the possibility that OA, like 5-HT, could influence the effects of 23E10 neurons on sleep and the FSB (Qian et al., 2017).

The expression patterns for the OA receptors we observed are in agreement with previous functional studies suggesting that OAMB and Octβ1R, but not Octβ2R, affect learning via expression in the MBs (Kim et al., 2013; Sabandal et al., 2020). Octopaminergic processes also innervate the EB, FSB and protocerebral bridge (PCB) (Sinakevitch and Strausfeld, 2006; Busch et al., 2009), which together with the noduli form the central complex. The potential functions of OAMB and Octβ1R in the central complex are not known. Our data suggest that genetic strategies to knockdown or knockout OAMB and Octβ1R in the central complex could be used to dissect the mechanisms by which this structure regulates behaviors such as orientation during flight (Seelig and Jayaraman, 2013; Green et al., 2017; Turner-Evans et al., 2017; Hardcastle et al., 2021), visually-evoked learning and memory (Pan et al., 2009), taste-independent nutrient selection (Dus et al., 2013), and sleep (Liu et al., 2016; Andreani et al., 2022; Kato et al., 2022; Yan et al., 2023).

### Octopamine receptors in the visual system

In addition to the MBs, octopaminergic neurons have been shown to innervate the optic lobe and regulate a wide range of visual behaviors (Monastirioti et al., 1995; Busch et al., 2009; Suver et al., 2012; Van Breugel et al., 2014; Cheng et al., 2019; Städele et al., 2020). However, the specific functions of different OA receptors in visual behavior remain unclear and their expression patterns in the visual system have not been described. In the optic lobe, we detect expression of OAMB-K3, Octβ1R and Octβ2R in several layers of the medulla, as well as the lobula and the lobula plate (Fig 3A-C). In contrast, we observe minimal detectable signal for OAMB-AS in the optic lobe (data not shown).

**Fig. 3:**
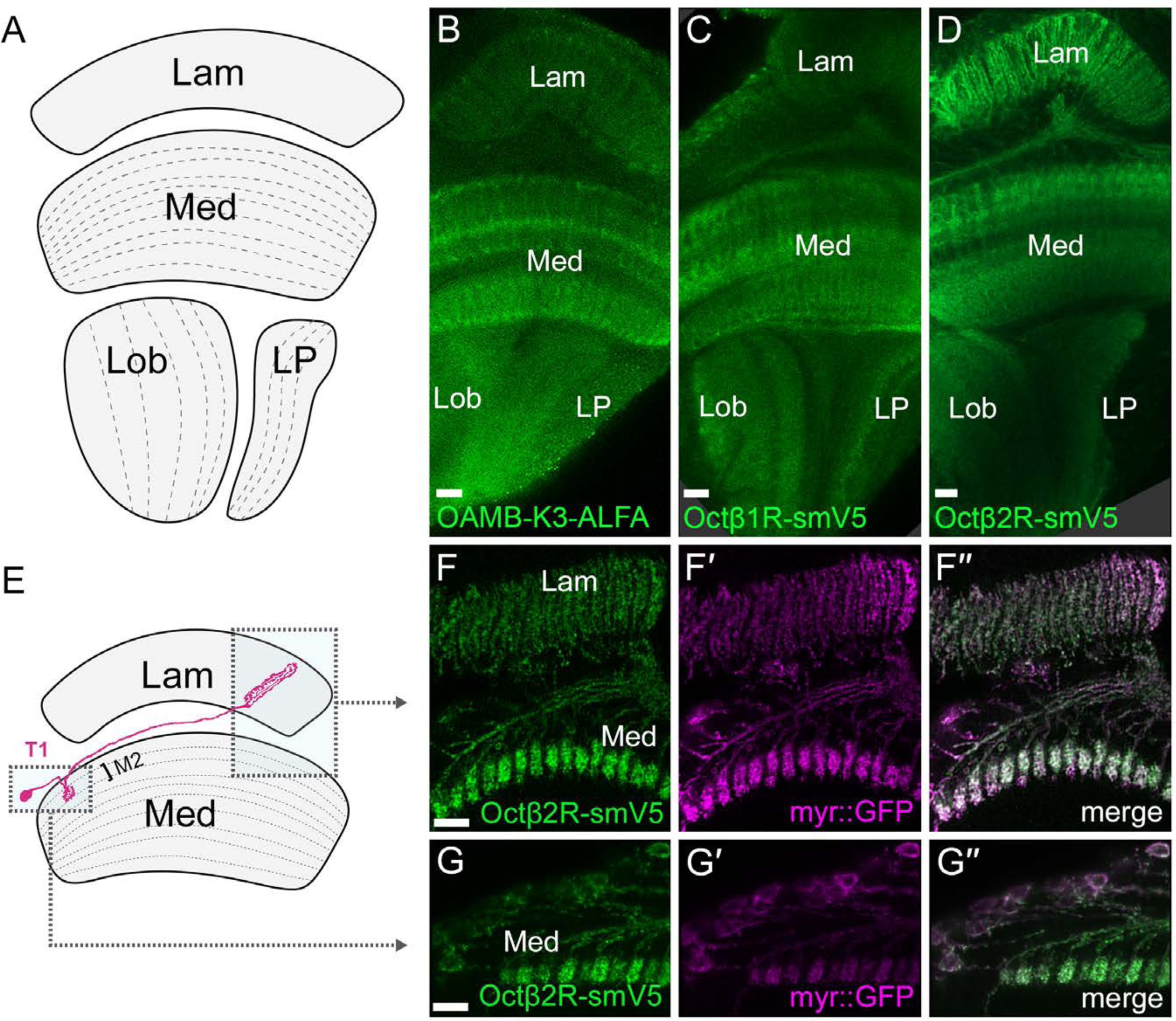
Octopamine receptors in the *Drosophila* optic lobe. A) Schematic of optic lobe showing each of the four major neuropils: lamina (Lam), medulla (Med), lobula (Lob) and lobula plate (LP). B) OAMB-K3 is expressed in at least two layers of the Med, but expressed at a level comparable to background in the Lam. B) Oct 1R is enriched in the Med, Lob and LP, but not the Lam. D) Oct 2R is expressed in the Med and the Lam. E) Schematic showing the regions of the Lam and Med imaged in F-G, including the projections of T1 cells (magenta) F-F′′) Conditional expression using a T1 specific driver of Oct 2R (green) and myr::GFP (magenta) confirms that labeled processes in the Med and Lam are derived from the T1 neuron. G-G′′) A higher resolution image of layer M2. Scale bars: 10μm.

Octβ2R-smV5 appears enriched in a distal (close to the eye) layer of the medulla that may be M2, and mRNA for the receptor is enriched in the morphologically distinct T1 cell that projects to M2 (Kurmangaliyev et al., 2020; Özel et al., 2021; Konstantinides et al., 2022). We thus hypothesized that T1 may be responsible for the Octβ2R signal in the distal medulla and to test this, we crossed a T1-specific driver (Tuthill et al., 2013) to the conditional receptor system. The data (Fig. 3D-D′′) confirm that Octβ2R is expressed in T1 and detectable in both the lamina and medulla neuropils (Fig. 3D-D′′, E-E′′). A large number of other cell-type-specific drivers are available for optic lobe neurons (Tuthill et al., 2013) and may be used to further analyze expression of OA receptors in the visual system.

In sum, OA receptors are enriched in multiple brain regions across the optic lobes and central brain. Using the tagged alleles to determine their localization to specific layers of the optic lobe neuropils may aid in the determination of their functions in visual behaviors.

### Serotonin receptors and the regulation of sleep and vision

Previous studies indicate that mutation of 5-HT1A results in a sleep deficit that can be rescued by specifically restoring expression in Kenyon cells (KCs) of the MBs (Yuan et al., 2006) (see cartoon, Fig. 4A), where it is also important in modulating the time-window for learning (Zeng et al., 2023). RNA-seq studies have also reported enrichment of 5-HT1A mRNA in one of the 3 major KC subtypes relative to the others (Aso et al., 2019; Bonanno and Krantz, 2023). We hypothesized that direct visualization of the receptor would mirror this enrichment and provide additional data regarding its subcellular localization. Indeed, we find that the constitutive allele of 5-HT1A is broadly expressed in the central brain and enriched in the MBs, particularly the α/β lobes (Fig. 4B). These results are also consistent with the distribution reported for a C-terminal sfGFP fusion of 5-HT1A (Deng et al., 2019).

**Fig. 4:**
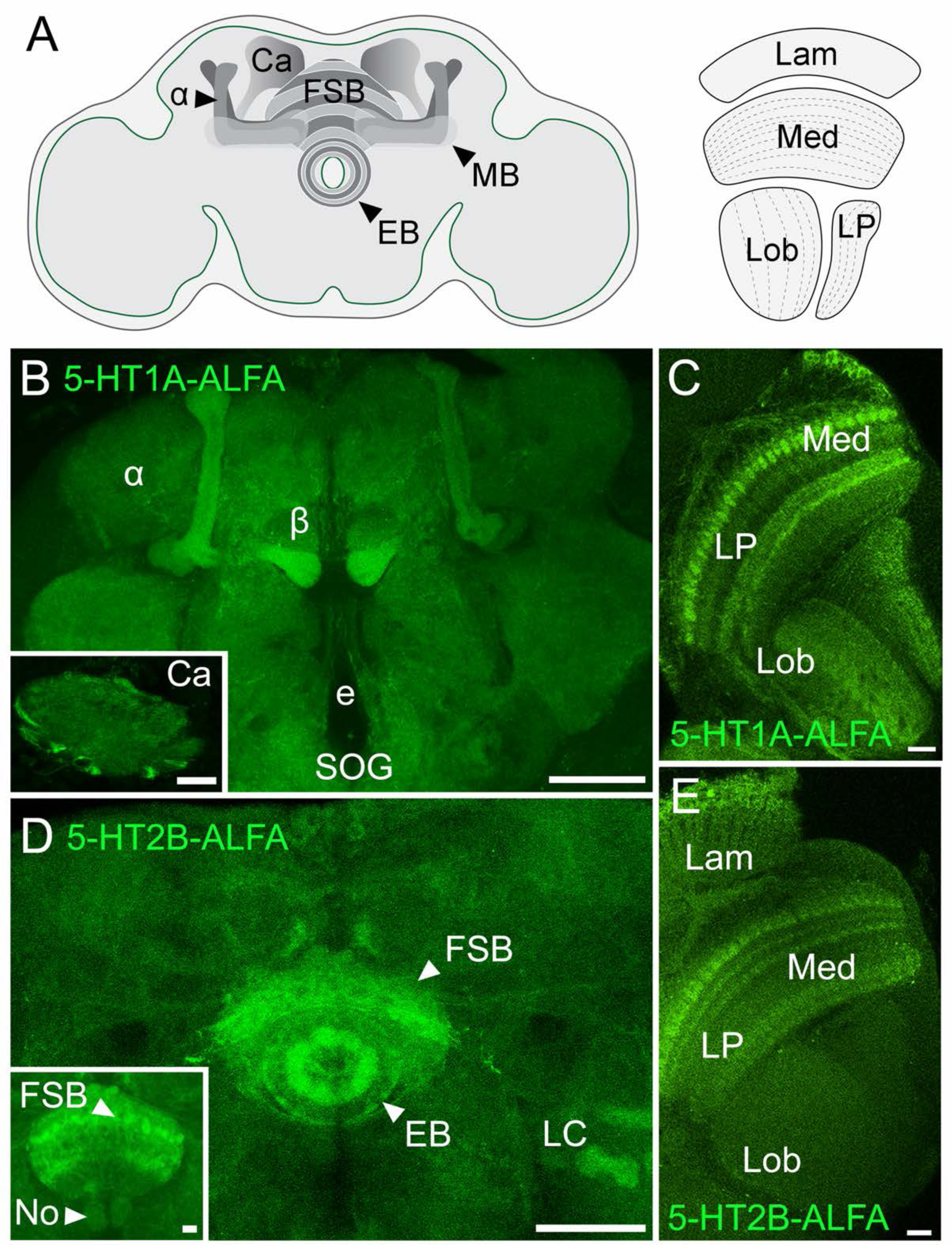
Serotonin receptors 5-HT1A and 5-HT2B localize to structures previously implicated in sleep and to the visual system. A) Schematic of the *Drosophila* brain, with the α lobe and calyx (Ca) of the mushroom body (MB), the ellipsoid body (EB) and fan shaped body (FSB) of the central complex, and the lamina (Lam), medulla (Med), lobula (Lob), and lobula plate (LP) of the optic lobe indicated. B) 5-HT1A is expressed throughout the central brain and enriched in the α/ lobes of the MB and the Ca (inset). e: esophagus; SOG: suboesophageal ganglion. C) In the optic lobe, 5-HT1A labels regions of the Med, Lob and LP. D) 5-HT2B is enriched in multiple layers of the FSB (see also inset) as well as the EB, the Noduli of the central complex (No) and at least 2 glomeruli of the lobular columnar cells (LC) that are visible here. E) 5-HT2B in the optic lobe is enriched in several layers of the Med as well as the Lam, but not detectable above background in the Lob or LP. Scale bars in B, D: 50μm; scale bars in C, E and in B, D insets: 10μm.

In the optic lobes, serotonergic innervation is broad (Nässel and Cantera, 1985; Vallés and White, 1988), and many cell types are enriched for one or more serotonin receptors (Kurmangaliyev et al., 2020; Sampson et al., 2020; Özel et al., 2021). We imaged 5-HT1A-ALFA in the optic lobes and observed expression in the medulla, lobula, and lobula plate, but none detectable above background in the lamina (Fig. 4C). Although the function of 5-HT1A in *Drosophila* vision is not known, its localization to the medulla could be the result of high 5-HT1A mRNA expression in T1 neurons which strongly innervate layer M2 (Kurmangaliyev et al., 2020; Özel et al., 2021).

Similar to 5-HT1A, 5-HT2B mutants also exhibit decreased sleep (Qian et al., 2017). While the 5-HT1A phenotype is mediated by expression in the MBs, 5-HT2B regulates sleep via activity in the FSB, and the mutant sleep phenotype can be rescued by restoration of receptor expression in the 23E10 neurons that project to the FSB (Qian et al., 2017). RNAi-mediated knockdown of 5-HT2B in 23E10 neurons also lead to social and grooming deficits (Cao et al., 2022), and previous work also revealed that 5-HT2B expressed by those cells traffics preferentially to their neuropil in the FSB and not their other, dorsal arborization (Qian et al., 2017).

Labeling of our 5-HT2B-ALFA allele showed that, consistent with previous studies, the receptor is enriched in several layers of the FSB, and additionally in the nearby ellipsoid body (EB) (Fig. 4D), but not the dorsal projections of 23E10. In the optic lobe, 5-HT2B-ALFA labeling in both distal and proximal layers (with respect to the central brain) of the medulla are prominent (Fig. 4E), while expression above background is not detectable in the lobula and lobula plate neuropils. The expression pattern of 5-HT2B in the distal layers of the medulla is consistent with previous transcriptomic data, as mRNA for the receptor is known to be enriched in L2 cells (Kurmangaliyev et al., 2020; Özel et al., 2021) which terminate in layer M2 (Bausenwein et al., 1992) and 5-HT2B is required for the response of L2 to serotonin (Sampson et al., 2020). In sum, although both 5-HT1A and 5-HT2B are crucial for the regulation of sleep and both receptors are expressed in a layer-specific fashion in the optic lobe, the cells that express each receptor and the neuropils to which they project are different, underscoring the complex serotonergic regulation of both sleep and visual behavior.

### The relationship of 5-HT2A to functional studies

To further explore the localization of 5-HT receptors in the central brain and optic lobe we imaged the constitutively tagged 5-HT2A allele. We observe a distinct labeling pattern for 5- HT2A-ALFA in the central brain (Fig. 5A-D) prompting us to review the influence of 5-HT2A on behavior and its potential relationship to specific anatomic structures. In the adult fly, loss of 5- HT2A leads to defects in feeding (Gasque et al., 2013; Munneke et al., 2022), increased lifespan (Munneke et al., 2022), and an altered response to the death of conspecifics (Chakraborty et al., 2019). The response to death has been linked to signaling in the EB, suggesting that 5-HT2A might be enriched at this site (Gendron et al., 2023). The 5-HT2A-ALFA signal we observe in the EB is above background, though more striking enrichment resides in bilateral foci at the distal edges of the FSB (Fig. 5B) which represent the lateral triangle, the first neuropil formed by the ring neurons whose projections form the EB (Hanesch et al., 1989; Young and Armstrong, 2010). Together these data suggest the possibility that 5-HT2A may preferentially localize to the somatodendritic compartment of cells that innervate the EB rather than axon terminals that innervate the FSB, and future experiments using the conditional allele will be used to test this hypothesis.

**Fig. 5:**
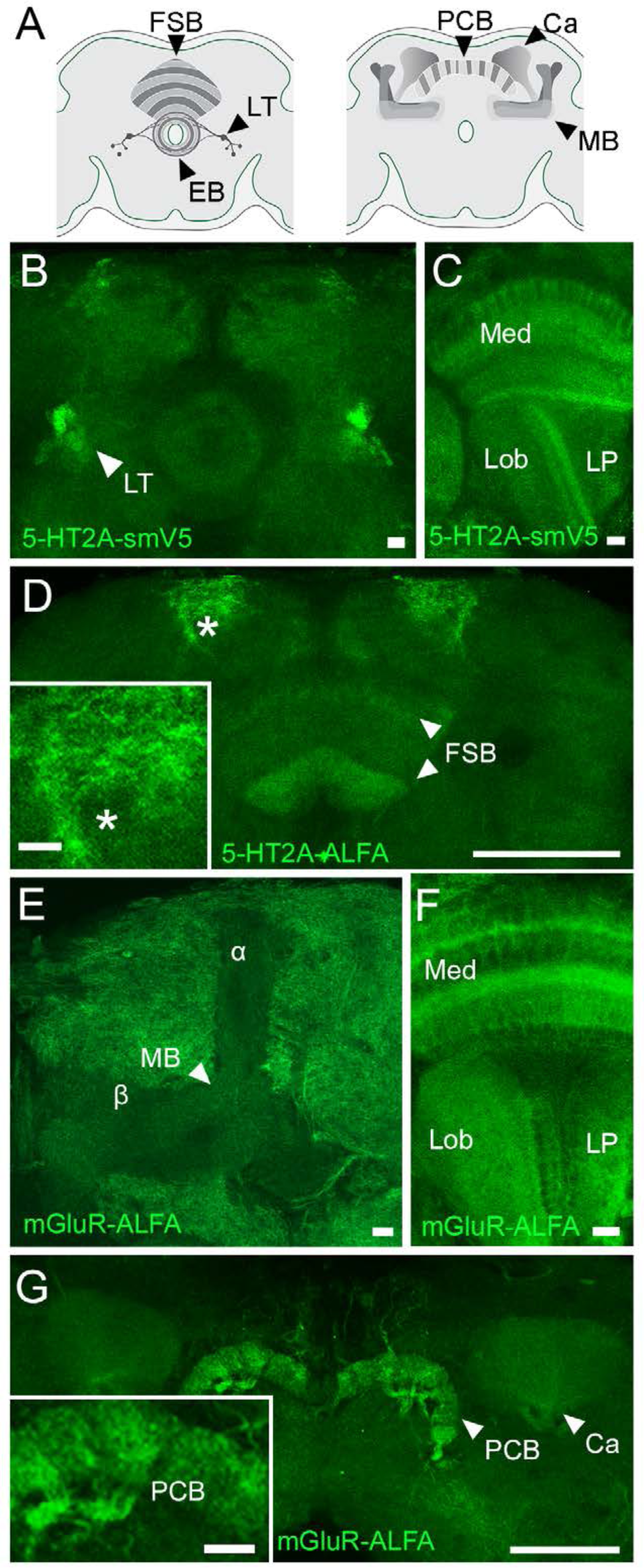
5-HT2A and mGluR are enriched in distinct neuropils in the central brain and optic lobe. A) Left: Schematic of structures labeled in the brain by 5-HT2A, including the fan-shaped body (FSB) and the ring neurons, which project into the lateral triangle (LT) and the ellipsoid body (EB). Right: Structures relevant to mGluR include the protocerebral bridge (PCB) and the mushroom bodies (MB) including the Calyx (Ca). B) 5-HT2A labels foci lateral to the EB that are morphologically identical to the lateral triangles (LT). C) In the optic lobe, 5-HT2A is expressed in the medulla (Med), and the lobula (Lob), with minimal signal in the lobula plate (LP). D) 5- HT2A is detected in ventral as well as a dorsal layers of the FSB, and highly expressed in a bilateral dorsal arborization (asterisk). E) mGluR is notably de-enriched from the lobes of the MB relative to the surrounding tissue. F) In the optic lobe, mGluR labeling is present in the Med, Lob and LP. G) mGluR is highly enriched in the protocerebral bridge (PCB) and also present in the Ca of the MBs. Scale bars in D,G: 50μm; all others:10μm.

We also detect labeling of the visual system with the tagged 5-HT2A allele, where the receptor is differentially enriched in distinct layers of the medulla, as well as in at least one layer of the lobula (Fig. 5C). Finally, 5-HT2A is also present in a bilateral arborization near the dorsal surface of the central brain and a dorsal as well as a ventral layer of the FSB (Fig. 5D).

### Expression of tagged mGluR in the MB lobes

While the primary focus of our studies was OA and 5-HT Class A GPCRs, mGluR provided an additional opportunity to test our ability to tag a class C GPCR. In addition, previously published data using an mGluR antibody provided us the opportunity to compare our labeling of tagged mGluR to a conventional antibody targeting the same protein. Functional studies suggest that expression of mGluR in the MBs facilitates learning (Schoenfeld et al., 2013; Andlauer et al., 2014) and its pharmacological activation can rescue behavioral deficits induced by mutation of the fragile X protein Fmr1 (McBride et al., 2005) as well as age-dependent sleep loss (Hou et al., 2023). We detect strong labeling of mGluR-ALFA in the PCB, as well as labeling in the MB calyces (Fig. 5E). By contrast, it appears to be de-enriched in the MB lobes relative to the surrounding tissue (Fig. 5F). This expression pattern is in accordance with previous data using the antibody raised against endogenous mGluR that is currently unavailable (Panneels et al., 2003; Bogdanik, 2004; Devaud et al., 2008) thereby confirming that the tag did not disrupt trafficking. We also observe mGluR-ALFA expression in the optic lobe, with enrichment in medial and distal layers of the medulla, as well as signal in both the lobula and lobula plate (Fig.5G).

Our observations on the 5-HT2A and mGluR tags may inform further studies of their function. In particular, the preferential labeling of 5-HT2A in the dendritic neuropil of putative ring neurons that form the lateral triangle compared to their axon terminals in the EB suggests the possibility that dendritic rather than axonal 5-HT2A receptors may be responsible for serotonergic modulation there. Similarly, it is possible that the primary site of activity of mGluR is the KC dendrites in the MB calyx rather than in the output neuropil that forms the lobes, given the low expression of mGluR in the MB lobes. This information is important because the subcellular site(s) of action for 5-HT2A, mGluR and, indeed most other GPCRs in the fly are not known. Their predicted localization to dendrites provides a testable hypothesis about the site at which they regulate neuronal activity.

### Expression of serotonin receptors in lobular columnar cells

Labeling the constitutive 5-HT2B-ALFA allele, we detect three neuropils corresponding to optic glomeruli whose morphology matches those of the terminals of particular lobula columnar (LC) cells (Wu et al., 2016): LC11, LC18, and LPLC2, with lower expression in the LC12 glomerulus that is more difficult to unambiguously distinguish from background (Fig. 6A,B). Although, we did not observe expression of 5-HT receptors other than 5-HT2B using the constitutive tags, we speculated other receptors may be expressed at lower levels and detected more easily using the conditional tagging system. We selected LC12 to test this hypothesis since our data on 5-HT2B already indicated that at least one 5-HT receptor was expressed at a relatively low level in these cells. To determine whether LC12 might express 5-HT1A and/or 5- HT2A in addition to 5-HT2B, we used our conditional alleles in combination with a split-GAL4 driver specific for LC12 (Wu et al., 2016). A myristoylated fluorophore was again used as a control to mark the membrane of labeled cells as for other labeling with the conditional alleles. We find that both 5-HT1A and 5-HT2A are expressed in LC12. Interestingly, 5-HT1A labeling appeared to be concentrated in the optic glomeruli relative to the dendritic neuropil in the lobula (Fig. 6C-D). These data raise the possibility that, like 5-HT1A in the lobes of the MBs, 5-HT1A may act to modulate neuronal activity at axonal sites within the optic LC neurons.

**Fig. 6:**
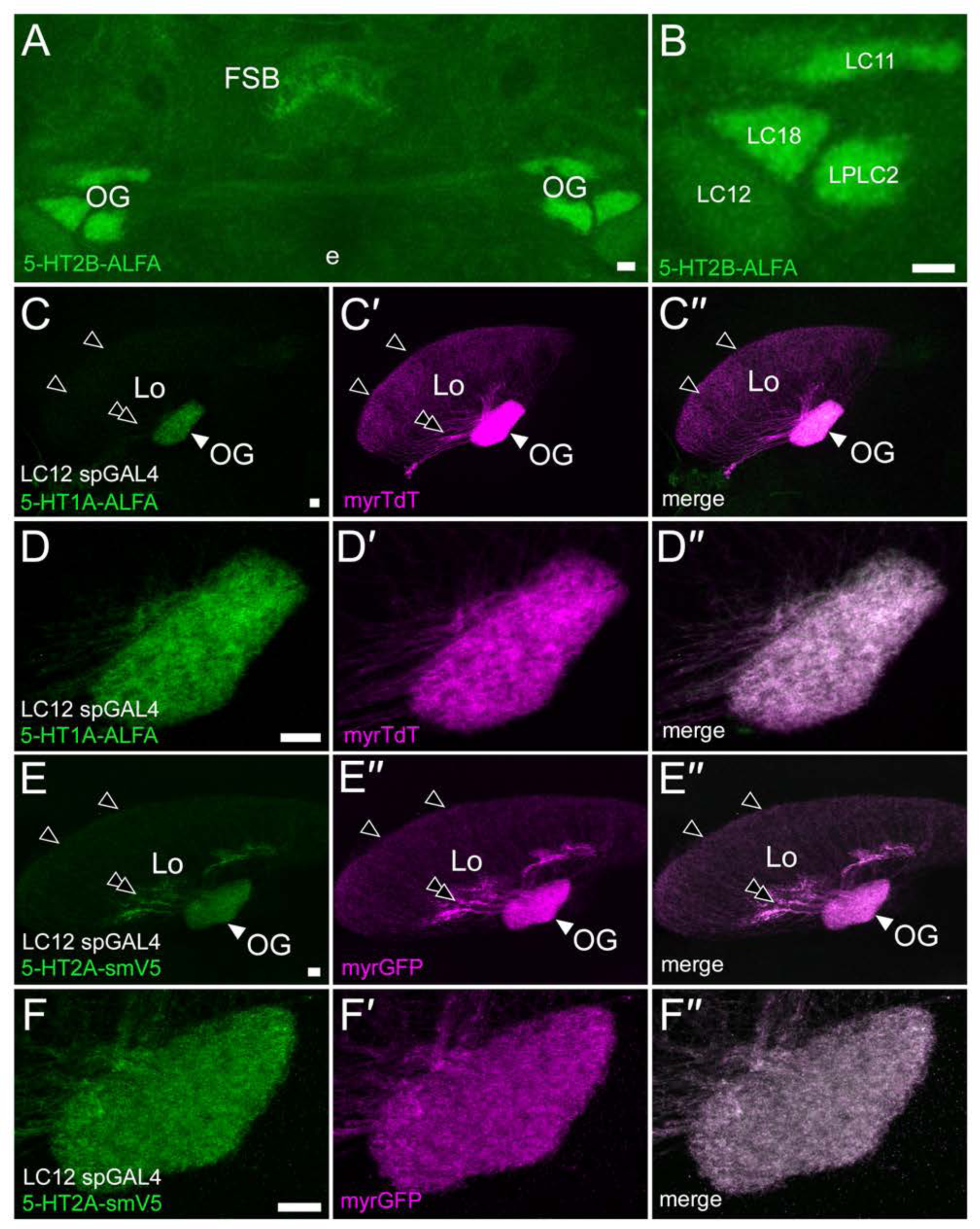
Expression of serotonin receptors in lobular columnar cells. A) Constitutive 5- HT2B-ALFA labeling shows enrichment in the FSB and optic glomeruli (OG). B) At higher magnification, the morphology of the OG suggest they likely represent projections from LC11, LC18, LPLC2 and LC12. C-C′′) Conditionally expressed 5HT1A-ALFA (green) is clearly present in the OG (white arrowhead), but more difficult to detect in either the proximal processes (double black arrowhead) or distal dendrites (single black arrowheads) within the lobula (Lo). Membrane bound TdTomato (magenta) is visible in both the OG and Lo. D-D′′) Higher magnification view of the LC12 OG. E-E′′) Conditionally expressed 5-HT2A is detectable in LC12. Labeling is clearly present in both the OG (white arrowhead) and proximal processes within the Lo (double black arrowhead); it is also detectable above background in the distal dendrites (single black arrowheads). E-E′′) High magnification view of 5-HT2A in the LC12 OG. Scale bars: 10μm.

### Comparison of presynaptic serotonin reuptake sites with postsynaptic receptors

Extrasynaptic serotonergic signaling and volume transmission have been proposed to play a fundamental role in the regulation of mammalian circuits (Descarries and Mechawar, 2000; Kaushalya et al., 2008; Vizi et al., 2010; Borroto-Escuela et al., 2015). In the absence of antibodies to map the location of 5-HT receptors, the distances between 5-HT release sites and their postsynaptic versus more distal targets in the fly brain have not been investigated. To facilitate the investigation of this question using the tagged 5-HT receptors, we generated an additional new marker for presynaptic serotonergic neurons using the *Drosophila* plasma membrane serotonin transporter (dSERT), (Lau et al., 2010; Kasture et al., 2019; Awasthi et al., 2021). We cloned N-terminally HA-tagged dSERT cDNA and inserted it into the MI^02578^ MiMIC site in the endogenous *Drosophila Sert* locus using recombinase-mediated cassette exchange (Fig. 7A) as previously described (Li-Kroeger et al., 2018). We then combined the tagged HA-SERT marker of presynaptic tracts with the constitutively tagged 5-HT1A and 5-HT2B alleles and performed co-labeling experiments.

**Fig. 7:**
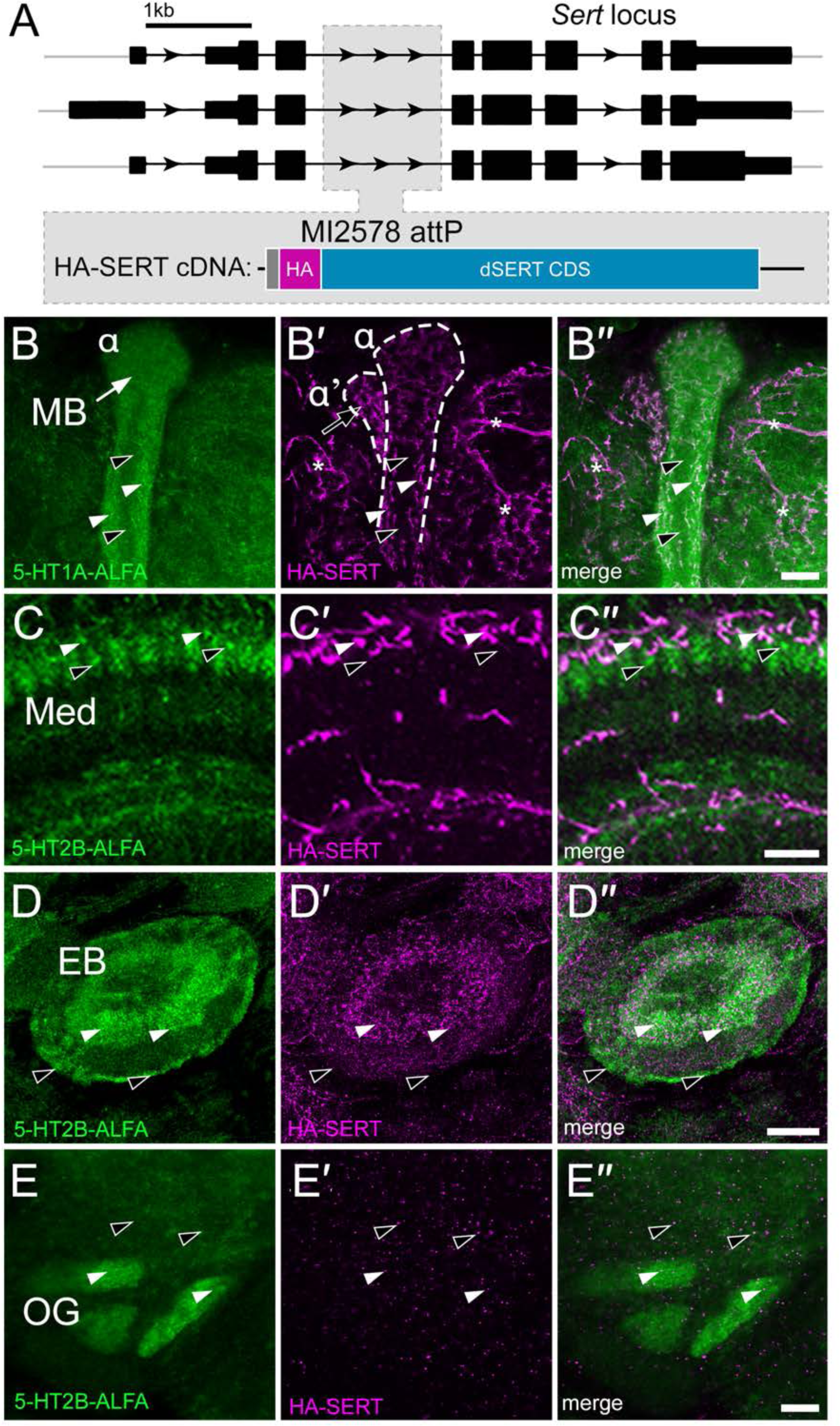
Comparison of presynaptic serotonin reuptake sites with serotonin receptors. A) Schematic of the *Drosophila Sert* (*dSERT*) genomic locus. The MiMIC insertion MI02578 between exons 3 and 4 in *dSERT* contains an attP site. The N-terminally HA-tagged *dSERT* coding sequence (CDS) was inserted into the attP site, preserving up- and downstream DNA regulatory regions. B-B′′) Constitutive 5-HT1A-ALFA (green) and HA-SERT (magenta) were co-labeled; the dorsal portion of α/α′ lobes are shown (B′, dotted white lines). 5-HT1A is enriched in α (B, white arrow) relative to α′ while the density of HA-SERT appears higher in α′ (B′, black arrow). Labeling of both 5-HT1A-ALFA and HA-SERT appear to be higher in the “shell” of α (B- B′′, white arrowheads), compared to the core (black arrowheads). Asterisks indicate HA-SERT labeling outside of the MB. C-C′′) The constitutive 5-HT2B-ALFA allele (green) was co-labeled with HA-SERT (magenta). The medulla (Med) shows areas in layer M2 that appear to co-label for 5-HT2B-ALFA and HA-SERT (C-C′′, white arrowheads) and others in which the receptor is detectable but HA-SERT is not (C-C′′, black arrowheads). D-D′′) 5-HT2B (green) and HA-SERT (magenta) were imaged together at the ellipsoid body (EB). Both the receptor and HA-SERT are enriched in the inner ring(s) of the EB (D-D′′, white arrowheads), but relatively little HA-SERT label is present in the outer ring (D-D′, black arrowheads). E-E′′) 5-HT2B (green) and HA-SERT (magenta) were imaged together in the region of the central brain that contains the optic glomeruli (OG) of the LC cells. Few HA-SERT(+) puncta were visible in this region either within the OG visible here (E-E′′, white arrowheads) or in the adjacent areas (E-E′′, black arrowheads). Scale bars: 10μm.

We first examined the spatial proximity of 5-HT1A-ALFA and HA-SERT signal in the MBs. We again observed enrichment of 5-HT1A-ALFA in the α lobe of the mushroom body (Fig. 7B), with serotonergic fibers in the outer shell as well as strong signal in areas surrounding the MB. (Fig. 7B′). The serotonergic fibers likely correspond to tracts originating from the serotonergic dorsal paired medial (DPM) and contralaterally projecting serotonin immunoreactive deutocerebral (CSD) cells, respectively (Pooryasin and Fiala, 2015; Scheffer et al., 2020). We also observe HA-SERT labeling in the α′ lobe (Fig. 7B′), although there is no corresponding enrichment of 5-HT1A in this region.

To extend our observations to other receptors and other brain regions, we examined the optic lobe, central complex, and optic glomeruli. 5-HT2B is expressed in the medulla of the optic lobe (Fig. 7C) and we have previously shown that when a tagged 5-HT2B transgene is expressed in laminar monopolar cell L2, it is enriched in layer M2 (Sampson et al., 2020). Interestingly, the serotonergic fibers labeled with HA-SERT appear to innervate a relatively limited portion of M2 that is closest to the surface of the eye (Fig. 7C′) and do not appear to be adjacent to the tagged receptors in other portions of this layer as would be expected for canonical, synaptic signaling. In the EB, we observe enrichment of 5-HT2B in both the inner and outer layers (Fig. 7D). While HA-SERT labeling appears proximal to 5-HT2B in the inner layer(s), we observe relatively fewer serotonergic fibers in the outer layer (Fig. 7D″). Finally, we imaged 5-HT2B and HA-SERT alleles in the area surrounding the optic glomeruli that house the terminals of LC neurons. Interestingly, though there is strong 5-HT2B signal in the LC glomeruli (Fig. 7D) we observed very sparse immunoreactivity corresponding to presynaptic serotonergic fibers (Fig. 7D′) as compared to the medulla of the optic lobe and the EB.

The surprising disconnect between the labeling of HA-SERT and 5-HT2B near the LC glomeruli raised the possibility that serotonin may be released at a site relatively distant from the target receptors. Although SERT has been consistently reported to mark the whole membrane of serotonergic cells (Lau et al., 2010; Kasture et al., 2019; Awasthi et al., 2021), it is possible that serotonin release might occur at sites unmarked by the reuptake transporter. To test this and to confirm that the HA-SERT labeling was indeed indicative of serotonergic innervation, we performed additional experiments to label the serotonergic processes within the region of the brain that houses the LC glomeruli. We crossed the broad serotonergic Tph-GAL4 driver (Park et al., 2006) to a line bearing UAS-mCD8::GFP and HA-SERT, such that the whole membrane of serotonergic cells would be marked by GFP and could be compared to the signal from the transporter. Surprisingly, though some of the LC glomeruli such as LC17 are innervated by serotonergic neurons, others such as the adjacent LC12 appear to be devoid of processes marked by HA-SERT or membrane-targeted GFP (Fig. 8A, B). To further investigate the serotonergic innervation of the LC12 and 17 glomeruli, we then plotted these neuron skeletons from the *Drosophila* hemibrain connectome (Scheffer et al., 2020) and included the 5-HTPLP and 5-HTPMP cells that provide most of the serotonergic innervation to the LC glomeruli (Pooryasin and Fiala, 2015; Scheffer et al., 2020). Consistent with the HA-SERT and Tph-GAL4>mCD8::GFP signal, the area innervated by LC17 is penetrated by serotonergic fibers while LC12 is not (Fig. 8C). While further experiments will be necessary, these data support the idea that the receptors in this region may be activated by 5-HT that diffuses from relatively distant terminals or is perhaps neurohumoral in origin.

**Fig. 8:**
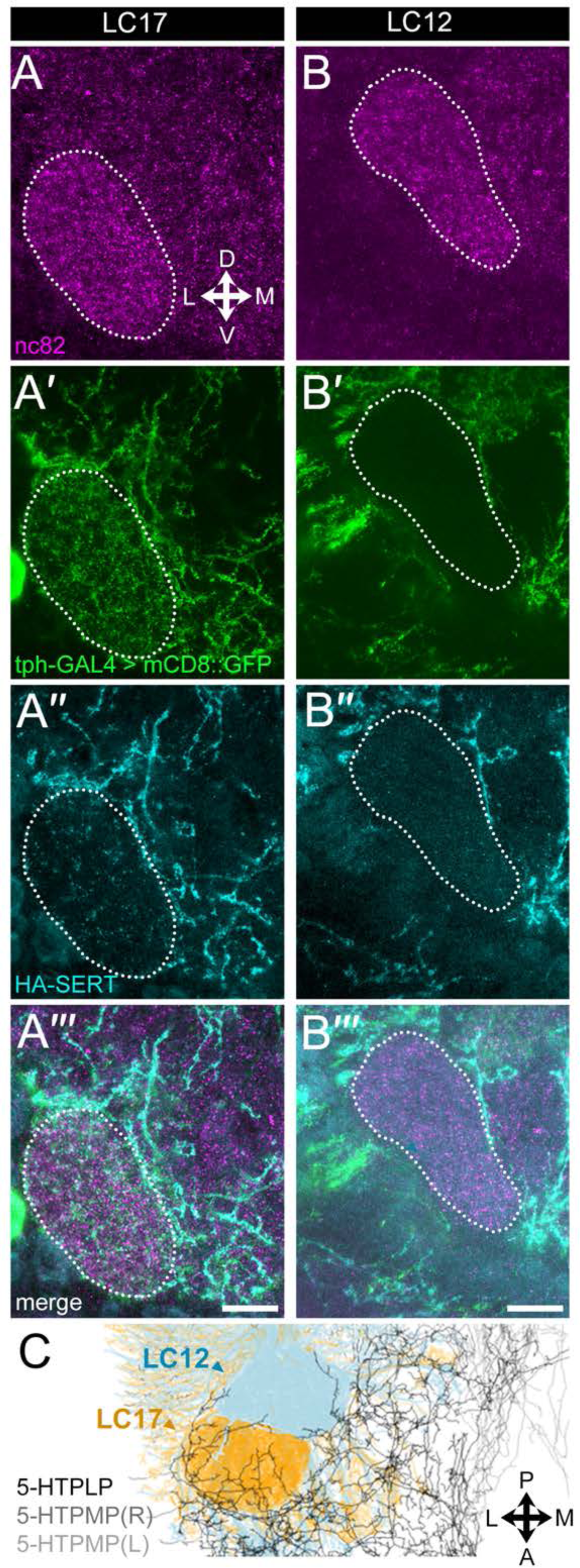
Serotonergic innervation of the LC glomeruli. All images are derived from the same confocal stack. A, B) nc82 (magenta) was used to label major neuropils and thus visualize the LC17 (A) and LC12 (B) glomeruli. A′, B′) The Tph-GAL4 driver was used to express membrane-targeted GFP and thereby label serotonergic processes (green). Serotonergic innervation of LC17 (A′) appears greater than LC12 (B′). A′′, B′′) HA-SERT (cyan) was used to co-label the serotonergic processes. Relatively few labeled puncta (white arrowheads) are present in LC12 compared to LC17. The intensity of A′′ and B′′ were altered to compensate for differences in background fluorescence. A′′′, B′′′) Merged images from panels A-A′′ and B-B′′. C) The *Janelia* hemibrain connectome was used to plot the morphology (skeletons) of the serotonergic neurons that innervate the LC glomeruli (5-HTPLP, 5-HTPMP(R), 5-HTPMP(L)), and the OG of both LC12 (light blue), and LC17 cells (orange). The field of view is rotated to top-down (dorso-ventral) so that both LCs can be visualized. Consistent with panels A-A′′ and B-B′′, fibers from serotonergic neurons appear to penetrate into the core of the OG of LC17 but not LC12. Scale bars: 10μm.

### Pre-versus postsynaptic expression of 5-HT1A

In mammals, some aminergic receptors are expressed in presynaptic nerve terminals as autoreceptors, which play a critical role in the regulation of neurotransmitter release (Richardson-Jones et al., 2010, 2011; Newman-Tancredi, 2011; Andrade et al., 2015; Milak et al., 2018). In *Drosophila,* mRNA studies suggest that aminergic receptors are also expressed in presynaptic aminergic neurons (Aso et al., 2019; Allen et al., 2020; Li et al., 2022) but their subcellular distribution and function is poorly understood. Even when available, using conventional antibodies or tagged receptors can make it difficult to discriminate between labeling of closely apposed proteins, which is a significant concern in tightly packed neuropils.

To explore the expression and localization of pre-versus postsynaptic 5-HT1A in the MBs we first used the *Janelia* fly connectome (Scheffer et al., 2020) to plot and quantify synapses to and from the main serotonergic neuron that provides input to the MB lobes, DPM (Fig. 9A-A′) which is both GABAergic and serotonergic, and synapses heavily onto all 3 of the major lobes of the mushroom body as well as accessory cells (Scheffer et al., 2020). RNA-seq data suggests that DPM expresses 5-HT1A mRNA (Aso et al., 2019) and its dense innervation of the MB indicates that the detection of 5-HT1A autoreceptors might be impeded by the high expression of postsynaptic 5-HT1A in the same region.

**Fig. 9:**
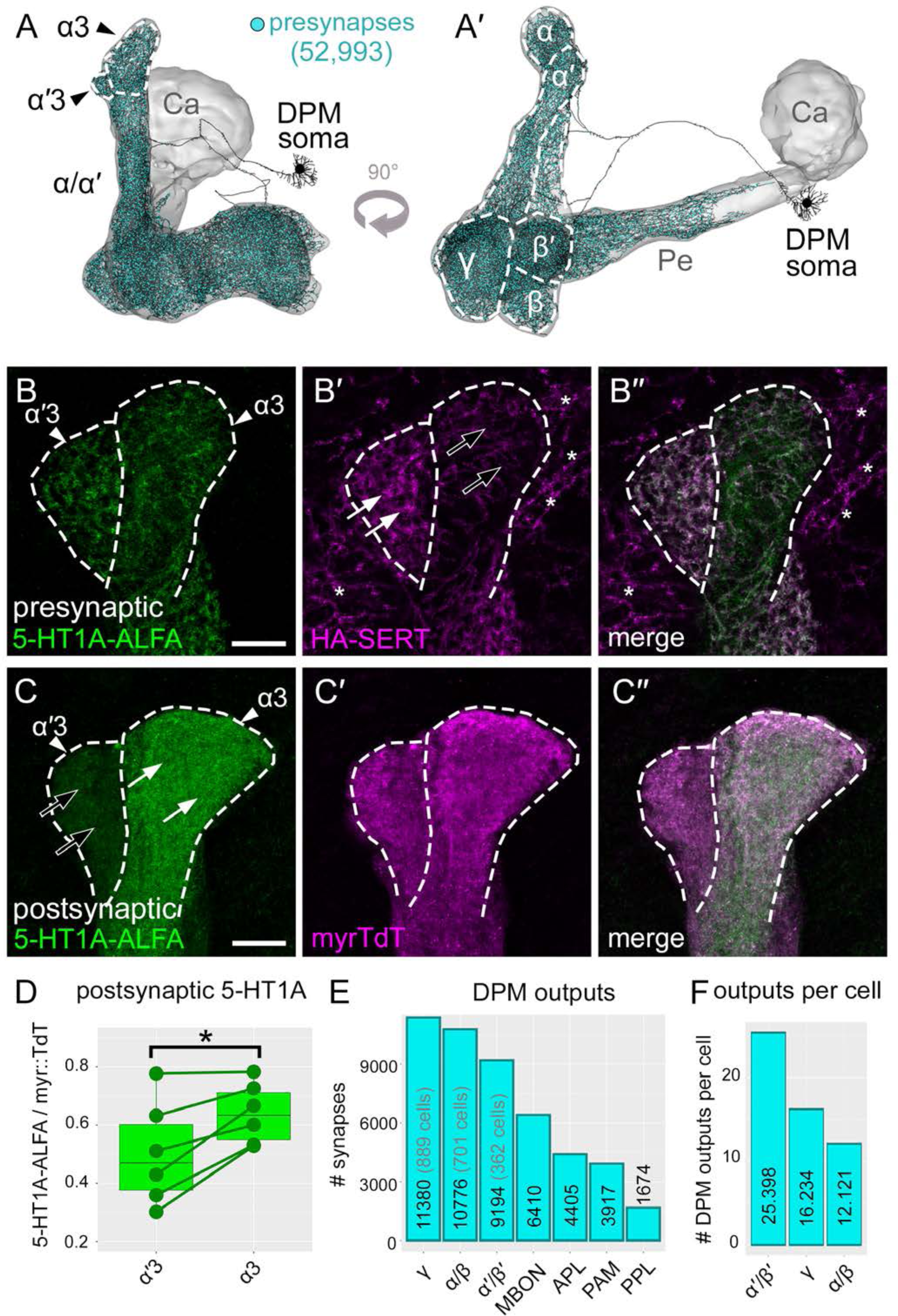
Conditional expression of 5-HT1A in pre-(DPM) and postsynaptic (KC) cells. A-A′) The anatomy of the serotonergic DPM neuron including the indicated soma and projections into the MB lobes (α/α′, β/β′ and γ) is shown in black, plotted as a “skeleton” from the neuprint connectome. The remaining portions of the mushroom body not innervated by DPM including the peduncle (Pe) and calyx (Ca) are shown in grey. The dorsal-most segments of the α/a′ lobes (α3 and α′3) are enclosed in dotted white lines here and in B and C below. B-B′′) The α3 and α′3 segments of the α/a′ lobes are shown presynaptically labeled with the conditional 5- HT1A-ALFA allele (green) expressed in DPM. HA-SERT co-label (magenta) was used to mark the presynaptic serotonergic membranes. Presynaptic 5-HT1A signal (B) is present in both α3 and α′3 and does not appear to be enriched in either lobe. HA-SERT (B′) appears enriched in α′3 (white arrows) compared to α3 (black arrows). Asterisks denote HA-SERT signal outside of the MB. C-C′′) The conditional 5-HT1A-ALFA allele (green) was expressed postsynaptically in KCs. 5-HT1A signal is clearly enriched in α3 (C, white arrows) relative to α′3 (C, black arrows). D) Quantification of the postsynaptic 5-HT1A receptor signal normalized to myr::TdT demonstrating the enrichment of 5-HT1A in α3 relative to α3′. p=0.01341 (paired t-test). E) Quantification of the number of DPM synapses onto each of the main cell types of the MB, using the connectome data plotted in (A). DPM primarily synapses onto KCs (γ, α/β, α′/β′) relative to other MB-extrinsic cells (MBONs, APL, PAMs, and PPLs). There are more synapses in total onto γ than other KC subtypes. F) Quantification of the number of DPM synapses as in E, but divided by the number of cells included in the analysis, per cell type (for each of the 3 KC classes). Though there are more synapses in total onto γ than other KC subtypes, the number of synapses per cell is highest on α′β′. Scale bars: 10μm.

To test the hypothesis that 5-HT1A is an autoreceptor in DPM and to determine its subcellular location, we used the DPM-specific c316-GAL4 driver in conjunction with the conditional receptor allele to restrict expression of 5-HT1A-ALFA. We co-labeled using the HA- SERT allele to compare the receptor localization with that of presynaptic 5-HT reuptake sites. 5- HT1A localizes to presynaptic processes of DPM within both α and α′ lobes, as does HA-SERT (Fig. 9B-B′′). To examine the postsynaptic expression of 5-HT1A in the same region, we crossed the conditional allele to OK107-GAL4 (Connolly et al., 1996), a pan-KC driver that expresses roughly evenly across α/β, α′/β′, and γ cells (Aso et al., 2009). Recapitulating transcriptional data (Aso et al., 2019; Bonanno and Krantz, 2023), 5-HT1A-ALFA is detectable in both α and α′ (Fig. 9C-C′′), but when normalized to myr:TdT in each lobe, it is enriched in α (Fig. 9D).

To further explore the distribution of DPM synapses onto each of the major cell types of the MB, we plotted its outputs using the *Janelia* connectome of the fly brain (Scheffer et al., 2020). DPM synapses heavily onto each of the 3 major KC subtypes, with γ receiving the most (Fig. 9E). When the total number of synapses for each type is normalized to the number of cells per type (# of DPM synapses per postsynaptic cell), however, the α′/ ′ cells that express the lowest levels of postsynaptic 5-HT1A receive the densest serotonergic innervation from DPM (Fig. 9F). These data reinforce the idea that the relationship between the sites of 5-HT release and its postsynaptic targets is complex and that the enrichment of presynaptic serotonergic fibers does not necessarily match the enrichment of any one specific postsynaptic receptor.

## Discussion

Key questions in the cell biology of neurons include the cellular and subcellular distribution of neuromodulatory receptors. Historically, the answers have relied on the generation of highly specific antibodies to receptors, but this has proven to be difficult for most *Drosophila* GPCRs. To address this issue, we exploited a strategy recently used for ionotropic receptors (Sanfilippo et al., 2023) to tag a number of GPCRs at their endogenous loci. The 5- HT1A, 5-HT2B, mGluR, and OAMB tagged alleles we have generated largely recapitulate published data on the enrichment of these receptors in particular brain regions (Han et al., 1998; Panneels et al., 2003; Bogdanik, 2004; Devaud et al., 2008; Kim et al., 2013; Deng et al., 2019). In lieu of custom antibodies that are either in limited supply or are no longer available, the tagged alleles render these targets once again accessible and introduce the potential for conditional labeling experiments. The localization of tagged receptors to the neuropil identified using endogenous antibodies to OAMB and mGluR indicate that tag placement in the 3^rd^ intracellular loop of Class A receptors and the extracellular domain of Class C GPCRs do not interfere significantly with receptor trafficking in *Drosophila*. Additionally, our success in tagging the two different protein isoforms of OAMB indicate that this strategy may be used to tag other GPCRs with alternative isoforms.

Data obtained using the tagged receptors has addressed several unanswered questions regarding their cellular expression and subcellular localization. One fundamental question is whether the receptor proteins are expressed in cells shown to express their mRNAs either in RNA-seq or transcriptional reporter studies, and we have indeed verified this for the subset of GPCRs that we have examined (Aso et al., 2019; Holt et al., 2019; Ament and Poulopoulos, 2023; Bonanno and Krantz, 2023). Importantly, previous efforts to define receptor-expressing cells that use GAL4 drivers to express broad cytosolic or plasma membrane markers do not necessarily match the expression patterns observed when directly visualizing the receptor. For example, the 5-HT2B-enriched LC glomeruli were not reported using GAL4 reporters (Gnerer et al., 2015), and the expression of Oct 1R in the 23E10 cells was similarly obscured by broad labeling of cell bodies and processes (McKinney et al., 2020). Moreover, while the use of GAL4 drivers and RNA-seq can reveal which cells may be enriched for particular receptors, they do not provide information on subcellular targeting – a key piece of information in neurons that often innervate multiple major neuropils and can be regulated at multiple subcellular sites including dendrites, axons and nerve terminals.

A second critical question in the study of neuromodulatory circuitry is the relationship between receptor expression and functional studies. We have confirmed the localization of 5- HT1A, 5-HT2B, OAMB, and Oct 1R proteins to the MBs and central complex in accordance with their demonstrated requirement in these structures for sleep and learning (Yuan et al., 2006; Modi et al., 2020; Zeng et al., 2023). In addition, we have generated several new hypotheses based on the expression patterns we observe. The enrichment of 5-HT2A and Oct 1R in specific layers of the FSB predicts the possibility of corresponding functional deficits in these circuits using receptor loss-of-function perturbations. We also observe enrichment of OAMB-K3 and Oct 1R in several different layers of the optic lobe neuropils. Although the optic lobes are innervated by octopaminergic neurons and OA has been shown to regulate multiple visual behaviors (Monastirioti et al., 1995; Aso et al., 2009; Suver et al., 2012; Cheng et al., 2019; Städele et al., 2020), the function of specific OA receptors in the fly visual system remains poorly defined. The enrichment of OA receptors in specific regions of the optic lobe that we show here may help to predict at which steps each one may regulate visual information processing.

The localization of receptors to specific subcellular domains that we observe generates additional hypotheses. In particular, the presence of 5-HT1A, 5-HT2A and 5-HT2B in the nerve terminals of the LC cells suggests the possibility that they may regulate local neurotransmitter release at these sites rather than or in addition to the dendritic neuropil in the lobula. Conversely, low expression of mGluR in the lobes of the MBs suggest that previously defined functions may rely on inputs to the dendritic field in the MB calyx. The spatial separation of axons and dendrites in both LCs and KCs highlights the feasibility of testing the effects of serotonergic inputs to each subcellular domain by manipulating different subsets of serotonergic neurons that target each region.

To further investigate the neuroanatomy of serotonergic innervation, we have also generated a new tagged allele of dSERT. We performed co-labeling experiments using HA- SERT with a subset of 5-HT receptors to explore the relationship between presynaptic serotonergic processes and their presumptive postsynaptic targets. Future experiments using higher resolution optical methods such as tissue expansion and super resolution microscopy will be needed to fully explore this relationship, though even at a gross anatomical level the mismatch between serotonergic innervation and receptor localization in some areas is striking. The apparent mismatch between presynaptic innervation and postsynaptic receptor is particularly pronounced for 5-HT1A, 5-HT2A, and 5-HT2B in the LC glomeruli. In the MB, a precedent for non-synaptic neuromodulatory transmission has been reported for dopamine with an estimated diffusional radius of ∼2 microns (Takemura et al., 2017). Together, these observations suggest that similar to many mammalian circuits (Agnati et al., 1992; Rice, 2000), some 5-HT receptors are relatively distant from serotonergic processes, and that volume rather than synaptic transmission may be the dominant mode of signaling at these sites. Recognizing this distinction will be crucial for understanding the mechanisms by which serotonin and other neuromodulatory amines regulate circuit activity and behavior since many molecular tools developed to study synaptic transmission cannot be applied to extra-synaptic signaling processes. In addition, for the vast majority of circuits and behaviors, the contribution of synaptic vs non-synaptic amine release is not known, and significant differences have been identified in the few instances when this has been explicitly examined (Grygoruk et al., 2014).

Finally, we used the conditionally tagged 5-HT1A receptor to separately visualize the pre- and postsynaptic receptor expression in the MB lobes and revealed different expression patterns for each. Identifying the location of aminergic autoreceptors has been critical for understanding their function in mammals and to determine whether they regulate global excitability via activities within dendrites or act at nerve terminals to regulate neurotransmitter release more locally (Richardson-Jones et al., 2010, 2011; Newman-Tancredi, 2011; Andrade et al., 2015; Milak et al., 2018). To our knowledge, this information has been essentially absent for *Drosophila* GPCRs, but is essential for understanding the functional consequences of mutating many aminergic receptors. For example, mutation of 5-HT1A has been shown to disrupt sleep in *Drosophila* (Yuan et al., 2006; Qian et al., 2017). While genetic rescue experiments indicate that postsynaptic expression in KCs can rescue some aspects of the phenotype (Yuan et al., 2006), our data raise the possibility that presynaptic expression of 5-HT1A in DPM could also contribute to changes in sleep observed in the 5-HT1A mutant, and perhaps other mutations that affect serotonin and sleep (Qian et al., 2017; Knapp et al., 2022).

The tagging strategy used in this study can be extended to other GPCRs including dopamine, tyramine, peptide, muscarinic acetylcholine, and GABA-B receptors, or other proteins involved in neuronal signaling. Similar tagging strategies have been used to tag some synaptic proteins such as vesicular transporters (Williams et al., 2019; Certel et al., 2022a, 2022b) and the components of postsynaptic structures (Parisi et al., 2023), though there are few examples of conditionally-tagged receptor alleles. Continued experiments using these tools to better understand the distribution of neuromodulatory receptors of different modalities across different cell types and brain circuits will be a key complement to ongoing efforts to map neuronal connectivity and the mechanisms by which circuits process information.

## Methods

### Identification of sites for insertion of tagging cassette in receptor sequences

*Drosophila melanogaster* GPCRs were chosen for tagging, and homologs from other *Drosophila* species as well as homologs/paralogs from more divergent insects, mouse, and human sequences were identified using the “Find similar sequences” function on UniProt KB (UniProt Consortium, 2023). Protein alignments were created using Clustal Omega (Sievers et al., 2011) and analyzed for regions that showed evidence of amino acid insertions/expansions (See Fig. 1A). Genomic DNA sequences corresponding to these regions were identified using the UCSC genome browser (Kent et al., 2002; Drosophila 12 Genomes Consortium et al., 2007; Raney et al., 2014; Hoskins et al., 2015), and a ∼2kb window surrounding the intended insertion site were downloaded using the Das server (Kent et al., 2002). Homology arms were then designed and cloned into a donor vector for homologous recombination as described below. An optimal targettable CRISPR/Cas9 site close to the intended site of genomic modification was identified using the UCSC genome browser, and assessed for predicted efficiency and specificity using the integrated measures (Moreno-Mateos et al., 2015; Doench et al., 2016).

### Molecular design of tagging cassette

The conditional tagging cassette in this study contains either 1xALFA tag (Götzke et al., 2019) or spaghetti-monster fluorescent protein with 10x V5 epitopes (smV5) (Viswanathan et al., 2015) flanked by 2xGGGGS linkers. The tag is preceded by a transcription interruption (STOP) cassette flanked by KD recombinase sites (KDRT) (Sanfilippo et al., 2023). The interruption cassette also includes a floxed 3xP3-DsRed selection marker, to assist in selection of transformants from embryo injections (see Fig. 1B).

### Molecular biology and cloning of vectors for tagged receptor alleles

Gibson Assembly (New England Biolabs, Cat# E2611S) or HiFi DNA Assembly (New England Biolabs, Cat# E5520S) was used to clone custom donor vectors for genomic integration as described in (Sanfilippo et al., 2023). DNA fragments for the 5′ and 3′ homology arms were obtained by PCR (∼1kb each), or by ordering gene fragments (∼150bp each) as described in (Kanca et al., 2019). Fragments corresponding to the tagging cassettes as described in (Sanfilippo et al., 2023) were digested from containing vectors. DNA fragments used in Gibson/HiFi assembly were gel purified, then sequentially PCR-purified using a commercial kit (Qiagen, Germantown, MD, Cat. No. 28506), and combined in an assembly reaction (NEB, Ipswitch, MA, Cat. No. 2621) to create a donor vector for genomic modification. Correct vector assembly was confirmed by colony-PCR, followed by Sanger sequencing. CRISPR sgRNA sequences were ordered as DNA oligos (IDT DNA, Coralville, IA), phosphorylated *in* vitro, and cloned into pU6-BbsI-gRNA (Addgene 45946, RRID:Addgene_45946) as previously described (Gratz et al., 2015). Detailed protocols are available upon request.

### Generation and verification of genetically-modified *Drosophila* lines

To make tagged receptor alleles, DNA for donor vectors and sgRNAs were sent to Best Gene, Inc. (Chino Hills, CA) for injection into *Drosophila* embryos from strains expressing Cas9 in the germline. G0 transformants were identified using the 3xP3-DsRed marker included in the tagging cassette, and crossed to balancers to create stable lines. These “founders” were crossed to Cyo-cre (Siegal and Hartl, 1996), to excise the floxed 3xP3-DsRed marker and create the final “conditional” allele, and then re-crossed to balancers to establish the allele in a *w^-^* background (see Fig. 1C). The same founders were crossed to a line expressing KDR in the germline (Sanfilippo et al., 2023) to excise all conditional portions of the tagging cassette and re-crossed to balancers yield the “constitutive” allele (see Fig. 1D).

To make the HA-SERT line, cDNA previously generated and cloned into pMT(HA-dSERT) (Romero-Calderón et al., 2007) was subcloned into the MiMIC RMCE vector: pBS-KS-attB1-2- PT-SA-SD-1 (DGRC plasmid 1305, RRID:DGRC_1305). This construct was injected into embryos from the SERT MiMIC MI02578 fly line at Best Gene, Inc., and transformants were identified and crossed to balancers to create stable lines.

### Fly husbandry

Flies were maintained on a standard cornmeal and molasses-based agar media with a 12:12 hour light/dark cycle at room temperature (22-25°C).

### Genetically-modified fly lines

The 5-HT1A-T2A-GAL4^MI01468^ fly line, described in (Gnerer et al., 2015) was a gift from Herman Dierick (Baylor College of Medicine).

The following fly lines were used in this study are as follows, with stock numbers for lines obtained from the Bloomington *Drosophila* Stock Center (BDSC, Bloomington, IN, USA) listed in parentheses. For those created in this study, they will be deposited to the Bloomington *Drosophila* Stock Center (BDSC, Bloomington, IN, USA):

UAS-mCD8::GFP (RRID:BDSC_5137)

Mef2(P247)-GAL4 (RRID:BDSC_50742)

3xUAS-FLPG5.PEST (RRID: BDSC_55808)

MB594-spGal4 (RRID:BDSC_69255)

T1 split-GAL4 (RRID:BDSC_68977, RRID:BDSC_70649)

### OK107-GAL4 (RRID:BDSC_854)

c316-GAL4 (RRID:BDSC_30830)

w, P{w+, 10xUAS-FRT-STOP-FRT-myrGFP-2A-KDR.PEST}attP40 (II) (gift from P. Sanfilippo, UCLA)

P{w+, 10xUAS-FRT-STOP-FRT-myrTdt-2A-KDR.PEST}attP5 (III) (gift from P. Sanfilippo, UCLA)

SerT MiMIC MI02578 (RRID:BDSC_36004)

### Created in this study

5-HT1A-K-ALFA (II)

5-HT1A-KSK-ALFA (II)

5-HT2A-K-smV5 (III)

5-HT2A-KSK-smV5 (III)

5-HT2B-K-ALFA (III)

5-HT2B-KSK-ALFA (III)

Oct 1R-K-ALFA (III)

Oct 1R-KSK-ALFA (III)

Oct 2R-K-ALFA (III)

Oct 2R-KSK-ALFA (III)

OAMB(K3)-K-ALFA (III)

OAMB(K3)-KSK-ALFA (III)

OAMB(AS)-K-ALFA (III)

OAMB(AS)-KSK-ALFA (III)

mGluR-K-ALFA (IV)

mGluR-KSK-ALFA (IV)

### Conditional receptor expression *in vivo*

For conditional receptor labeling experiments, fly lines were built and crossed such that progeny bore the conditional receptor allele plus three additional transgenes: a GAL4 driving cell-type restricted expression of the transcription factor, a UAS-FLP to drive FLP recombinase in the GAL4-defined population, and the 10xUAS-FRT-STOP-FRT-myrTdTom-T2A-KDR (or 10xUAS- FRT-STOP-FRT-myrGFP-T2A-KDR). In all cells except those expressing GAL4, the conditional receptor allele retains its KDRT-dependent stop cassette, and there is no production of epitope-tagged receptor. In GAL4-expressing cells, FLP excises the STOP cassette in the 10xUAS- FRT-STOP-FRT-myrTdTom-T2A-KDR transgene, permitting expression of myristoylated fluorophore and KDR (as independent transcripts). KDR then excises the STOP cassette in the conditional receptor, thus permitting both membrane-labeled fluorophore and epitope-tagged receptor in the GAL4-defined population.

### Immunofluorescent labeling and imaging

Adult female fly brains were dissected in ice-cold Schneider′s Media (ThermoFisher, Cat#21720024, Waltham, MA), then transferred to Terasaki multiwell plates (Millipore Sigma, Cat#M5812, St. Louis, MO) for the rest of the immunohistochemistry protocol. Brains were fixed in 3% glyoxal with acetic acid (Sigma-Aldrich, Cat#128465, St. Louis, MO) for 25 minutes at room temperature. After fixation, brains were washed three times in PBS with 0.5% Triton X-100 (Millipore Sigma, Cat#X100, Burlington, MA) (PBST) for 10 minutes, then blocked for 60 minutes in PBST containing 5% normal goat serum (NGS) (Cayman Chemical, Cat#10006577, Ann Arbor, MA). Solution was then switched to 5% NGS/PBST containing primary antibodies (see below) overnight for two days at 4°C. Brains were then washed by switching solution to PBST three times 60 minutes, then incubated with secondary antibodies overnight for two days in the dark at 4°C. Finally, brains were washed three times with PBST for 30 minutes, equilibrated in 50% Vectashield mounting media (Vector Labs, Cat#H-1900, Burlingame, CA) for 15 minutes, and then 100% mounting media for 10 minutes before transferring to slides and mounting with a coverslip. For imaging the central brain, whole brains were mounted horizontally (anterio-posterior). To image the visual system, tissue was mounted vertically (dorso-ventral) to capture all four of the major neuropils: lamina, medulla, lobula, and lobula plate (Fischbach and Dittrich, 1989).

### Primary antibodies

Myristoylated GFP (myr::GFP) was labeled with 1:500 mouse anti-GFP (Sigma-Aldrich, Cat#G6539, RRID:AB_259941, or ThermoFisher, Cat#A-11120, RRID:AB_221568)

Myristoylated TdTomato (myr::TdTom) was labeled with 1:500 rabbit anti-DsRed (Takara Cat# 632496, RRID:AB_10013483)

HA was labeled with 1:500 rabbit anti-HA (Cell Signaling Technology, Cat#3724, Danvers, MA, RRID:AB_1549585), 1:500 mouse anti-HA (BioLegend, Cat#MMS-101P, RRID_AB_2314672, San Diego, USA), or 1:500 rat anti-HA (Roche, Cat#1215817001, RRID:AB_390915, Indianapolis, USA)

smV5 was labeled 1:500 mouse anti-V5 (Thermo Fisher, Cat#MA5-15253, RRID:AB_10977225).

ALFA was labeled with 1:500 mouse Fc-conjugated nanobody anti-ALFA (NanoTag Biotechnologies, Cat#N1582, RRID:AB_3065196 Göttingen, Germany)

Brp was labeled with mouse nc82 1:100 (Developmental Studies Hybridoma Bank, Cat#nc82, RRID: AB_2314866, Iowa City, USA)

Secondary antibodies were used at 1:500 and include: Goat anti-mouse Alexa Fluor 488 (ThermoFisher, Cat#A32723, RRID:AB_2633275), Goat anti-mouse Alexa Fluor 568 (ThermoFisher, Cat#A11004, RRID:AB_2534072), Goat anti-Rabbit Alexa Fluor 488 (ThermoFisher, Cat#A11008, RRID:AB_143165), and Goat anti-Rabbit Alexa Fluor 555 (ThermoFisher, Cat#A21428, RRID:AB_2535849).

Imaging was performed with a Zeiss LSM 880 Confocal with Airyscan (Zeiss, Oberkochen, Germany) using a 20x air, 40x glycerol, or 63x oil immersion objective. Post-hoc processing of images was done with Fiji (Schindelin et al., 2012) or Adobe Photoshop (Adobe, San Jose, CA).

### Quantification of confocal images

For quantification in Figure 9D, flies bearing the 5-HT1A conditional receptor allele and conditional transgenes as well as Mef2-GAL4 were dissected and brains were stained with anti-ALFA (5-HT1A) and DsRed (myr::TdT) antibodies. Z-stacks (n=6) were acquired encompassing the volume of both α and α′, then imported to Fiji and z-projections were generated using maximum signal intensity. The myr::TdT channel was used to draw a mask around the α and α′ lobes, then average signal intensity for both ALFA and myr::TdT was measured in each region individually and recorded. The signal for ALFA was then normalized to myr::TdT per region, then data points were plotted as paired observations (i.e. normalized ALFA signal in α and α′ within one image) and statistical significance in difference between the means was assessed using a paired t-test in R (version 4.3.1). This analysis utilized the R packages ggplot2 (Wickham, 2016) and tidyverse (Wickham et al., 2019).

### Connectome analysis

The neuprintr R package (Bates et al., 2023) was used to plot DPM (body id: 5813105172) from the hemibrain:v1.2.1 construction of the *Janelia* fly connectome, extract data on synapse number and identity, and was further analyzed using standard data processing methods in R.

## Funding

This work was funded by HHMI (SLZ) an NIH BRAIN initiative award (1RF1MH117823-01) (SLZ and DEK) and R01MH114017 (DEK). The funders had no role in study design, data collection and analysis, decision to publish, or preparation of the manuscript.

## Contributions

PS and SLZ designed and developed the tagging system. SLB and DEK conceived of and designed all experiments. AE and MMS created HA-dSERT. BK and KL assisted in design and cloning for creation of tagged receptors. GDB performed molecular (PCR) validation of new fly lines created in this study. SLB performed all immunohistochemistry and imaging. SLB, DEK, PS and SLZ wrote the manuscript.

## Conflicts of interests

There are no competing financial interests in relation to the work described.

## Conflict of interest statement

The authors declare no competing financial interests.

## Acknowledgments

This work was funded by HHMI (SLZ) an NIH BRAIN initiative award (1RF1MH117823-01) (SLZ and DEK) and R01MH114017 (DEK).We would like to acknowledge Mark Dombrovskiy, Alex Kim, Juyoun Yoo, Saumya Jain, and other members of the Zipursky Lab for helpful discussion on experiments and antibody selection.

---

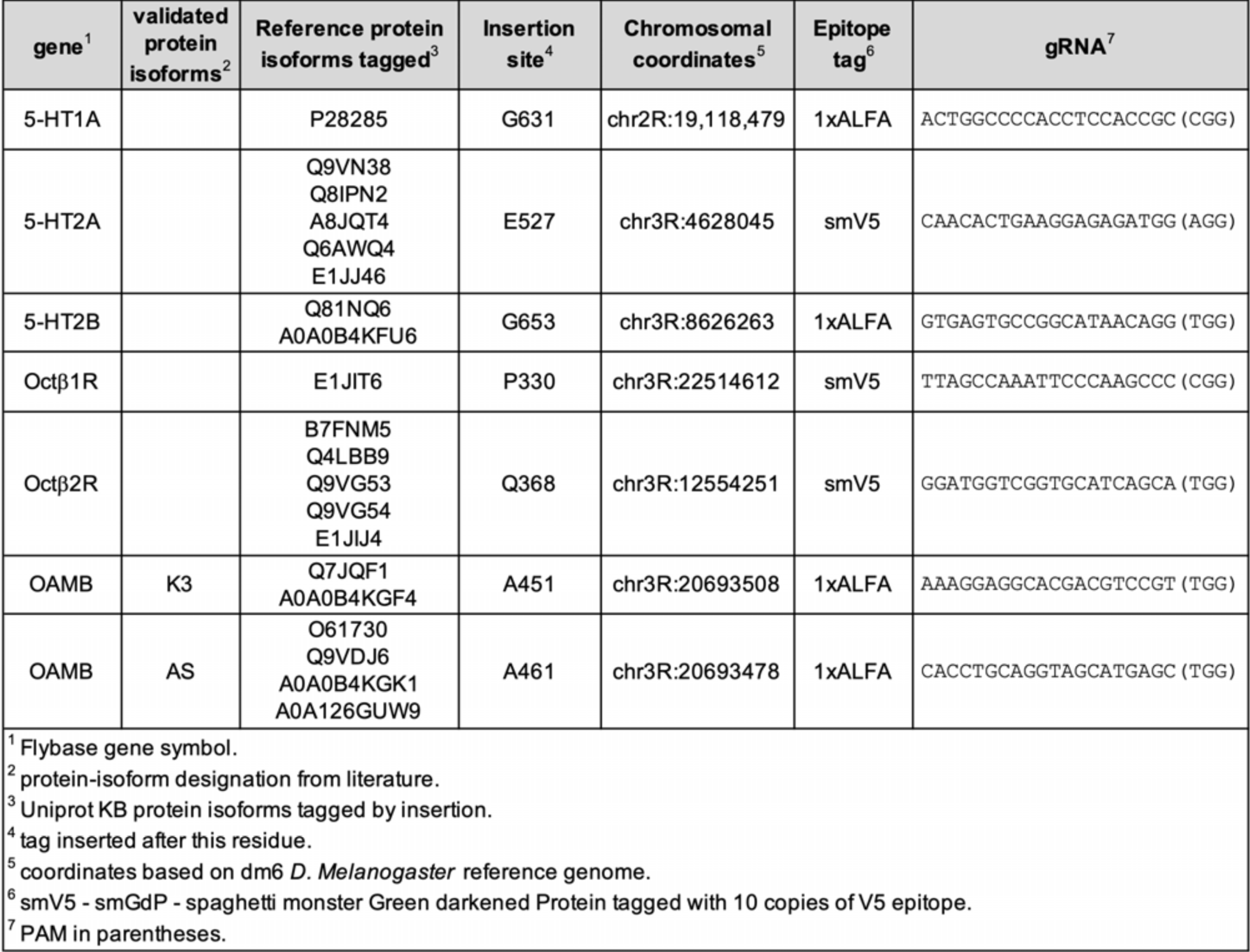

## Notes

### Competing Interest Statement

The authors have declared no competing interest.

## References

Agnati LF, Bjelke B, Fuxe K (1992) Volume Transmission in the Brain. American Scientist 80:362–373.

Akalal D-BG, Wilson CF, Zong L, Tanaka NK, Ito K, Davis RL (2006) Roles for Drosophila mushroom body neurons in olfactory learning and memory. Learn Mem 13:659–668.

Albert PR, Vahid-Ansari F (2019) The 5-HT1A receptor: Signaling to behavior. Biochimie 161:34–45.

Alekseyenko OV, Chan Y-B, Okaty BW, Chang Y, Dymecki SM, Kravitz EA (2019) Serotonergic Modulation of Aggression in Drosophila Involves GABAergic and Cholinergic Opposing Pathways. Current Biology 29:2145–2156.e5.

Allen AM, Neville MC, Birtles S, Croset V, Treiber CD, Waddell S, Goodwin SF (2020) A single-cell transcriptomic atlas of the adult Drosophila ventral nerve cord Mann RS, VijayRaghavan K, Sims PA, eds. eLife 9:e54074.

Ament SA, Poulopoulos A (2023) The brain’s dark transcriptome: Sequencing RNA in distal compartments of neurons and glia. Current Opinion in Neurobiology 81:102725.

Andersson H, D’Antona AM, Kendall DA, Von Heijne G, Chin C-N (2003) Membrane assembly of the cannabinoid receptor 1: impact of a long N-terminal tail. Mol Pharmacol 64:570– 577.

Andlauer TFM, Scholz-Kornehl S, Tian R, Kirchner M, Babikir HA, Depner H, Loll B, Quentin C, Gupta VK, Holt MG, Dipt S, Cressy M, Wahl MC, Fiala A, Selbach M, Schwärzel M, Sigrist SJ (2014) Drep-2 is a novel synaptic protein important for learning and memory. eLife 3:e03895.

Andrade R, Huereca D, Lyons JG, Andrade EM, McGregor KM (2015) 5-HT1A Receptor-Mediated Autoinhibition and the Control of Serotonergic Cell Firing. ACS Chem Neurosci 6:1110–1115.

Andreani T, Rosensweig C, Sisobhan S, Ogunlana E, Kath W, Allada R (2022) Circadian programming of the ellipsoid body sleep homeostat in Drosophila. eLife 11:e74327.

Aso Y, Grübel K, Busch S, Friedrich AB, Siwanowicz I, Tanimoto H (2009) The Mushroom Body of Adult *Drosophila* Characterized by GAL4 Drivers. Journal of Neurogenetics 23:156– 172.

Aso Y, Ray RP, Long X, Bushey D, Cichewicz K, Ngo T-T, Sharp B, Christoforou C, Hu A, Lemire AL, Tillberg P, Hirsh J, Litwin-Kumar A, Rubin GM (2019) Nitric oxide acts as a cotransmitter in a subset of dopaminergic neurons to diversify memory dynamics. eLife 8:e49257.

Awasthi JR, Tamada K, Overton ETN, Takumi T (2021) Comprehensive topographical map of the serotonergic fibers in the male mouse brain. Journal of Comparative Neurology 529:1391–1429.

Bates A, Jefferis G, Franconville R (2023) neuprintr: R client utilities for interacting with the neuPrint connectome analysis service. Available at: https://natverse.org/neuprintr.

Bausenwein B, Dittrich APM, Fischbach K-F (1992) The optic lobe of Drosophila melanogaster. Cell Tissue Res 267:17–28.

Bogdanik L (2004) The Drosophila Metabotropic Glutamate Receptor DmGluRA Regulates Activity-Dependent Synaptic Facilitation and Fine Synaptic Morphology. Journal of Neuroscience 24:9105–9116.

Bonanno SL, Krantz DE (2023) Transcriptional changes in specific subsets of Drosophila neurons following inhibition of the serotonin transporter. Transl Psychiatry 13:226.

Borroto-Escuela DO, Agnati LF, Bechter K, Jansson A, Tarakanov AO, Fuxe K (2015) The role of transmitter diffusion and flow versus extracellular vesicles in volume transmission in the brain neural–glial networks. Philosophical Transactions of the Royal Society B: Biological Sciences 370:20140183.

Busch S, Selcho M, Ito K, Tanimoto H (2009) A map of octopaminergic neurons in the Drosophila brain. Journal of Comparative Neurology 513:643–667.

Cao H, Tang J, Liu Q, Huang J, Xu R (2022) Autism-like behaviors regulated by the serotonin receptor 5-HT2B in the dorsal fan-shaped body neurons of Drosophila melanogaster. European Journal of Medical Research 27:203.

Certel SJ, McCabe BD, Stowers RS (2022a) A conditional GABAergic synaptic vesicle marker for Drosophila. Journal of Neuroscience Methods 372:109540.

Certel SJ, Ruchti E, McCabe BD, Stowers RS (2022b) A conditional glutamatergic synaptic vesicle marker for Drosophila. G3 (Bethesda) 12:jkab453.

Chakraborty TS, Gendron CM, Lyu Y, Munneke AS, DeMarco MN, Hoisington ZW, Pletcher SD (2019) Sensory perception of dead conspecifics induces aversive cues and modulates lifespan through serotonin in Drosophila. Nat Commun 10:2365.

Cheng KY, Colbath RA, Frye MA (2019) Olfactory and Neuromodulatory Signals Reverse Visual Object Avoidance to Approach in Drosophila. Current Biology 29:2058–2065.e2.

Cho D, Min C, Jung K, Cheong S, Zheng M, Cheong S, Oak M, Cheong J, Lee B, Kim K (2012) The N-terminal region of the dopamine D2 receptor, a rhodopsin-like GPCR, regulates correct integration into the plasma membrane and endocytic routes. British Journal of Pharmacology 166:659–675.

Connolly JB, Roberts IJ, Armstrong JD, Kaiser K, Forte M, Tully T, O’Kane CJ (1996) Associative learning disrupted by impaired Gs signaling in Drosophila mushroom bodies. Science 274:2104–2107.

Del-Bel E, De-Miguel FF (2018) Extrasynaptic Neurotransmission Mediated by Exocytosis and Diffusive Release of Transmitter Substances. Frontiers in Synaptic Neuroscience 10.

Deng B, Li Q, Liu X, Cao Y, Li B, Qian Y, Xu R, Mao R, Zhou E, Zhang W, Huang J, Rao Y (2019) Chemoconnectomics: Mapping Chemical Transmission in Drosophila. Neuron 101:876–893.e4.

Descarries L, Mechawar N (2000) Ultrastructural evidence for diffuse transmission by monoamine and acetylcholine neurons of the central nervous system. In: Progress in Brain Research, pp 27–47 Volume Transmission Revisited. Elsevier.

Devaud J-M, Clouet-Redt C, Bockaert J, Grau Y, Parmentier M-L (2008) Widespread brain distribution of the Drosophila metabotropic glutamate receptor. Neuroreport 19:367–371.

Donlea JM, Pimentel D, Miesenböck G (2014) Neuronal Machinery of Sleep Homeostasis in Drosophila. Neuron 81:860–872.

Donlea JM, Pimentel D, Talbot CB, Kempf A, Omoto JJ, Hartenstein V, Miesenböck G (2018) Recurrent Circuitry for Balancing Sleep Need and Sleep. Neuron 97:378–389.e4.

Dunham JH, Hall RA (2009) Enhancement of G Protein-Coupled Receptor Surface Expression. Trends Biotechnol 27:541–545.

Dus M, Ai M, Suh GSB (2013) Taste-independent nutrient selection is mediated by a brain-specific Na+/solute co-transporter in Drosophila. Nat Neurosci 16:526–528.

Feinberg EH, VanHoven MK, Bendesky A, Wang G, Fetter RD, Shen K, Bargmann CI (2008) GFP Reconstitution Across Synaptic Partners (GRASP) Defines Cell Contacts and Synapses in Living Nervous Systems. Neuron 57:353–363.

Fendl S, Vieira RM, Borst A (2020) Conditional protein tagging methods reveal highly specific subcellular distribution of ion channels in motion-sensing neurons. eLife 9:e62953.

Fischbach K-F, Dittrich APM (1989) The optic lobe of Drosophila melanogaster. I. A Golgi analysis of wild-type structure. Cell Tissue Res 258:441–475.

Fuxe K, Borroto-Escuela D, Romero-Fernandez W, Diaz Cabiale Z, Rivera A, Ferraro L, Tanganelli S, Tarakanov A, Garriga P, Narvaez JA, Ciruela F, Agnati L (2012) Extrasynaptic Neurotransmission in the Modulation of Brain Function. Focus on the Striatal Neuronal–Glial Networks. Frontiers in Physiology 3A.

Garcia-Garcia A, Tancredi AN-, Leonardo ED (2014) 5-HT1A receptors in mood and anxiety: recent insights into autoreceptor versus heteroreceptor function. Psychopharmacology (Berl) 231:623–636.

Gasque G, Conway S, Huang J, Rao Y, Vosshall LB (2013) Small molecule drug screening in Drosophila identifies the 5HT2A receptor as a feeding modulation target. Sci Rep 3:srep02120.

Gendron CM, Chakraborty TS, Duran C, Dono T, Pletcher SD (2023) Ring neurons in the Drosophila central complex act as a rheostat for sensory modulation of aging. PLOS Biology 21:e3002149.

Gnerer JP, Venken KJT, Dierick HA (2015) Gene-specific cell labeling using MiMIC transposons. Nucleic Acids Res 43:e56.

Götzke H, Kilisch M, Martínez-Carranza M, Sograte-Idrissi S, Rajavel A, Schlichthaerle T, Engels N, Jungmann R, Stenmark P, Opazo F, Frey S (2019) The ALFA-tag is a highly versatile tool for nanobody-based bioscience applications. Nat Commun 10:4403.

Gratz SJ, Rubinstein CD, Harrison MM, Wildonger J, O’Connor-Giles KM (2015) CRISPR-Cas9 genome editing in Drosophila. Curr Protoc Mol Biol 111:31.2.1-31.2.20.

Green J, Adachi A, Shah KK, Hirokawa JD, Magani PS, Maimon G (2017) A neural circuit architecture for angular integration in Drosophila. Nature 546:101–106.

Grygoruk A, Chen A, Martin CA, Lawal HO, Fei H, Gutierrez G, Biedermann T, Najibi R, Hadi R, Chouhan AK, Murphy NP, Schweizer FE, Macleod GT, Maidment NT, Krantz DE (2014) The Redistribution of Drosophila Vesicular Monoamine Transporter Mutants from Synaptic Vesicles to Large Dense-Core Vesicles Impairs Amine-Dependent Behaviors. J Neurosci 34:6924–6937.

Guan XM, Kobilka TS, Kobilka BK (1992) Enhancement of membrane insertion and function in a type IIIb membrane protein following introduction of a cleavable signal peptide. Journal of Biological Chemistry 267:21995–21998.

Han K-A, Millar NS, Davis RL (1998) A Novel Octopamine Receptor with Preferential Expression inDrosophila Mushroom Bodies. J Neurosci 18:3650–3658.

Hanesch U, Fischbach K-F, Heisenberg M (1989) Neuronal architecture of the central complex in Drosophila melanogaster. Cell Tissue Res 257:343–366.

Hardcastle BJ, Omoto JJ, Kandimalla P, Nguyen B-CM, Keleş MF, Boyd NK, Hartenstein V, Frye MA (2021) A visual pathway for skylight polarization processing in Drosophila Desplan C, Calabrese RL, Heinze S, Franconville R, eds. eLife 10:e63225.

Holt CE, Martin KC, Schuman EM (2019) Local translation in neurons: visualization and function. Nature Structural & Molecular Biology 26:557–566.

Hou X, Hayashi R, Itoh M, Tonoki A (2023) Small-molecule screening in aged Drosophila identifies mGluR as a regulator of age-related sleep impairment. Sleep 46:zsad018.

Kanca O et al. (2019) An efficient CRISPR-based strategy to insert small and large fragments of DNA using short homology arms VijayRaghavan K, VijayRaghavan K, eds. eLife 8:e51539.

Kasture AS, Bartel D, Steinkellner T, Sucic S, Hummel T, Freissmuth M (2019) Distinct contribution of axonal and somatodendritic serotonin transporters in drosophila olfaction. Neuropharmacology 161:107564.

Kato YS, Tomita J, Kume K (2022) Interneurons of fan-shaped body promote arousal in Drosophila. PLoS One 17:e0277918.

Kaushalya SK, Nag S, Ghosh H, Arumugam S, Maiti S (2008) A high-resolution large area serotonin map of a live rat brain section. NeuroReport 19:717.

Kim Y-C, Lee H-G, Lim J, Han K-A (2013) Appetitive Learning Requires the Alpha1-Like Octopamine Receptor OAMB in the Drosophila Mushroom Body Neurons. J Neurosci 33:1672–1677.

Knapp EM, Kaiser A, Arnold RC, Sampson MM, Ruppert M, Xu L, Anderson MI, Bonanno SL, Scholz H, Donlea JM, Krantz DE (2022) Mutation of the Drosophila melanogaster serotonin transporter dSERT impacts sleep, courtship, and feeding behaviors. PLOS Genetics 18:e1010289.

Kondo S, Takahashi T, Yamagata N, Imanishi Y, Katow H, Hiramatsu S, Lynn K, Abe A, Kumaraswamy A, Tanimoto H (2020) Neurochemical Organization of the Drosophila Brain Visualized by Endogenously Tagged Neurotransmitter Receptors. Cell Reports 30:284–297.e5.

Konstantinides N, Holguera I, Rossi AM, Escobar A, Dudragne L, Chen Y-C, Tran TN, Martinez Jaimes AM, Özel MN, Simon F, Shao Z, Tsankova NM, Fullard JF, Walldorf U, Roussos P, Desplan C (2022) A complete temporal transcription factor series in the fly visual system. Nature 604:316–322.

Kurmangaliyev YZ, Yoo J, Valdes-Aleman J, Sanfilippo P, Zipursky SL (2020) Transcriptional Programs of Circuit Assembly in the Drosophila Visual System. Neuron 108:1045–1057.e6.

Lau T, Schneidt T, Heimann F, Gundelfinger ED, Schloss P (2010) Somatodendritic serotonin release and re-uptake in mouse embryonic stem cell-derived serotonergic neurons. Neurochemistry International 57:969–978.

Li H et al. (2022) Fly Cell Atlas: A single-nucleus transcriptomic atlas of the adult fruit fly. Science 375:eabk2432.

Li-Kroeger D, Kanca O, Lee P-T, Cowan S, Lee MT, Jaiswal M, Salazar JL, He Y, Zuo Z, Bellen HJ (2018) An expanded toolkit for gene tagging based on MiMIC and scarless CRISPR tagging in Drosophila Yamashita YM, VijayRaghavan K, eds. eLife 7:e38709.

Liu S, Liu Q, Tabuchi M, Wu MN (2016) Sleep Drive Is Encoded by Neural Plastic Changes in a Dedicated Circuit. Cell 165:1347–1360.

Macpherson LJ, Zaharieva EE, Kearney PJ, Alpert MH, Lin T-Y, Turan Z, Lee C-H, Gallio M (2015) Dynamic labelling of neural connections in multiple colours by trans-synaptic fluorescence complementation. Nat Commun 6:10024.

Maqueira B, Chatwin H, Evans PD (2005) Identification and characterization of a novel family of Drosophilaβ-adrenergic-like octopamine G-protein coupled receptors. Journal of Neurochemistry 94:547–560.

McBride SMJ, Choi CH, Wang Y, Liebelt D, Braunstein E, Ferreiro D, Sehgal A, Siwicki KK, Dockendorff TC, Nguyen HT, McDonald TV, Jongens TA (2005) Pharmacological Rescue of Synaptic Plasticity, Courtship Behavior, and Mushroom Body Defects in a Drosophila Model of Fragile X Syndrome. Neuron 45:753–764.

McDonald NA, Henstridge CM, Connolly CN, Irving AJ (2007) An Essential Role for Constitutive Endocytosis, but Not Activity, in the Axonal Targeting of the CB1 Cannabinoid Receptor. Mol Pharmacol 71:976–984.

McKinney HM, Sherer LM, Williams JL, Certel SJ, Stowers RS (2020) Characterization of Drosophila octopamine receptor neuronal expression using MiMIC-converted Gal4 lines. Journal of Comparative Neurology 528:2174–2194.

Milak MS, Pantazatos S, Rashid R, Zanderigo F, DeLorenzo C, Hesselgrave N, Ogden RT, Oquendo MA, Mulhern ST, Miller JM, Burke AK, Parsey RV, Mann JJ (2018) Higher 5- HT1A autoreceptor binding as an endophenotype for major depressive disorder identified in high risk offspring – A pilot study. Psychiatry Research: Neuroimaging 276:15–23.

Modi MN, Shuai Y, Turner GC (2020) The Drosophila Mushroom Body: From Architecture to Algorithm in a Learning Circuit. Annual Review of Neuroscience 43:465–484.

Monastirioti M, Gorczyca M, Rapus J, Eckert M, White K, Budnik V (1995) Octopamine immunoreactivity in the fruit fly Drosophila melanogaster. Journal of Comparative Neurology 356:275–287.

Munneke AS, Chakraborty TS, Porter SS, Gendron CM, Pletcher SD (2022) The serotonin receptor 5-HT2A modulates lifespan and protein feeding in Drosophila melanogaster. Frontiers in Aging 3.

Nässel DR, Cantera R (1985) Mapping of serotonin-immunoreactive neurons in the larval nervous system of the flies Calliphora erythrocephala and Sarcophaga bullata. Cell Tissue Res 239:423–434.

Newman-Tancredi A (2011) Biased agonism at serotonin 5-HT 1A receptors: preferential postsynaptic activity for improved therapy of CNS disorders. Neuropsychiatry 1:149– 164.

Özel MN, Simon F, Jafari S, Holguera I, Chen Y-C, Benhra N, El-Danaf RN, Kapuralin K, Malin JA, Konstantinides N, Desplan C (2021) Neuronal diversity and convergence in a visual system developmental atlas. Nature 589:88–95.

Pan Y, Zhou Y, Guo C, Gong H, Gong Z, Liu L (2009) Differential roles of the fan-shaped body and the ellipsoid body in Drosophila visual pattern memory. Learn Mem 16:289–295.

Panneels V, Eroglu C, Cronet P, Sinning I (2003) Pharmacological characterization and immunoaffinity purification of metabotropic glutamate receptor from Drosophila overexpressed in Sf9 cells. Protein Expression and Purification 30:275–282.

Parisi MJ, Aimino MA, Mosca TJ (2023) A conditional strategy for cell-type-specific labeling of endogenous excitatory synapses in Drosophila. Cell Reports Methods 3:100477.

Park J, Lee SB, Lee S, Kim Y, Song S, Kim S, Bae E, Kim J, Shong M, Kim J-M, Chung J (2006) Mitochondrial dysfunction in Drosophila PINK1 mutants is complemented by parkin. Nature 441:1157–1161.

Pimentel D, Donlea JM, Talbot CB, Song SM, Thurston AJF, Miesenböck G (2016) Operation of a homeostatic sleep switch. Nature 536:333–337.

Pooryasin A, Fiala A (2015) Identified Serotonin-Releasing Neurons Induce Behavioral Quiescence and Suppress Mating in Drosophila. J Neurosci 35:12792–12812.

Qian Y, Cao Y, Deng B, Yang G, Li J, Xu R, zhang D, Huang J, Rao Y (2017) Sleep homeostasis regulated by 5HT2b receptor in a small subset of neurons in the dorsal fan-shaped body of drosophila Sehgal A, ed. eLife 6:e26519.

Rice ME (2000) Distinct regional differences in dopamine-mediated volume transmission. In: Progress in Brain Research, pp 277–290 Volume Transmission Revisited. Elsevier.

Richardson-Jones JW, Craige CP, Guiard BP, Stephen A, Metzger KL, Kung HF, Gardier AM, Dranovsky A, David DJ, Beck SG, Hen R, Leonardo ED (2010) 5-HT1A Autoreceptor Levels Determine Vulnerability to Stress and Response to Antidepressants. Neuron 65:40–52.

Richardson-Jones JW, Craige CP, Nguyen TH, Kung HF, Gardier AM, Dranovsky A, David DJ, Guiard BP, Beck SG, Hen R, Leonardo ED (2011) Serotonin-1A Autoreceptors Are Necessary and Sufficient for the Normal Formation of Circuits Underlying Innate Anxiety. J Neurosci 31:6008–6018.

Roeder T (2005) Tyramine and Octopamine: Ruling Behavior and Metabolism. Annual Review of Entomology 50:447–477.

Romero-Calderón R, Shome RM, Simon AF, Daniels RW, DiAntonio A, Krantz DE (2007) A screen for neurotransmitter transporters expressed in the visual system of Drosophila melanogaster identifies three novel genes. Dev Neurobiol 67:550–569.

Sabandal JM, Sabandal PR, Kim Y-C, Han K-A (2020) Concerted Actions of Octopamine and Dopamine Receptors Drive Olfactory Learning. J Neurosci 40:4240–4250.

Sampson MM, Myers Gschweng KM, Hardcastle BJ, Bonanno SL, Sizemore TR, Arnold RC, Gao F, Dacks AM, Frye MA, Krantz DE (2020) Serotonergic modulation of visual neurons in Drosophila melanogaster Desplan C, ed. PLoS Genet 16:e1009003.

Sanfilippo P, Kim AJ, Bhukel A, Yoo J, Mirshahidi PS, Pandey V, Bevir H, Yuen A, Mirshahidi PS, Guo P, Li H-S, Wohlschlegel JA, Aso Y, Zipursky SL (2023) Mapping of multiple neurotransmitter receptor subtypes and distinct protein complexes to the connectome.:2023.10.02.560011 Available at: https://www.biorxiv.org/content/10.1101/2023.10.02.560011v1.

Scheffer LK et al. (2020) A connectome and analysis of the adult Drosophila central brain Marder E, Eisen MB, Pipkin J, Doe CQ, eds. eLife 9:e57443.

Schindelin J, Arganda-Carreras I, Frise E, Kaynig V, Longair M, Pietzsch T, Preibisch S, Rueden C, Saalfeld S, Schmid B, Tinevez J-Y, White DJ, Hartenstein V, Eliceiri K, Tomancak P, Cardona A (2012) Fiji: an open-source platform for biological-image analysis. Nat Methods 9:676–682.

Schoenfeld BP, Choi RJ, Choi CH, Terlizzi AM, Hinchey P, Kollaros M, Ferrick NJ, Koenigsberg E, Ferreiro D, Leibelt DA, Siegel SJ, Bell AJ, McDonald TV, Jongens TA, McBride SMJ (2013) The Drosophila DmGluRA is required for social interaction and memory. Front Pharmacol 4:64.

Seelig JD, Jayaraman V (2013) Feature detection and orientation tuning in the Drosophila central complex. Nature 503:262–266.

Shearin HK, Quinn CD, Mackin RD, Macdonald IS, Stowers RS (2018) t-GRASP, a targeted GRASP for assessing neuronal connectivity. J Neurosci Methods 306:94–102.

Siegal ML, Hartl DL (1996) Transgene Coplacement and High Efficiency Site-Specific Recombination with the Cre/Loxp System in Drosophila. Genetics 144:715–726.

Sinakevitch I, Strausfeld NJ (2006) Comparison of octopamine-like immunoreactivity in the brains of the fruit fly and blow fly. Journal of Comparative Neurology 494:460–475.

Sorkaç A, Moșneanu RA, Crown AM, Savaş D, Okoro AM, Memiş E, Talay M, Barnea G (2023) retro-Tango enables versatile retrograde circuit tracing in Drosophila Hiesinger PR, Desplan C, Hiesinger PR, Luo L, eds. eLife 12:e85041.

Städele C, Keleş MF, Mongeau J-M, Frye MA (2020) Non-canonical receptive field properties and neuromodulation of feature detecting neurons in flies. Curr Biol 30:2508–2519.e6.

Suver MP, Mamiya A, Dickinson MH (2012) Octopamine Neurons Mediate Flight-Induced Modulation of Visual Processing in Drosophila. Current Biology 22:2294–2302.

Takemura S et al. (2017) A connectome of a learning and memory center in the adult Drosophila brain Calabrese RL, ed. eLife 6:e26975.

Turner-Evans D, Wegener S, Rouault H, Franconville R, Wolff T, Seelig JD, Druckmann S, Jayaraman V (2017) Angular velocity integration in a fly heading circuit. eLife 6:e23496.

Tuthill JC, Nern A, Holtz SL, Rubin GM, Reiser MB (2013) Contributions of the 12 Neuron Classes in the Fly Lamina to Motion Vision. Neuron 79:128–140.

Vallés AM, White K (1988) Serotonin-containing neurons in Drosophila melanogaster: Development and distribution. Journal of Comparative Neurology 268:414–428.

Van Breugel F, Suver MP, Dickinson MH (2014) Octopaminergic modulation of the visual flight speed regulator of *Drosophila*. Journal of Experimental Biology:jeb.098665.

Viswanathan S, Williams ME, Bloss EB, Stasevich TJ, Speer CM, Nern A, Pfeiffer BD, Hooks BM, Li W-P, English BP, Tian T, Henry GL, Macklin JJ, Patel R, Gerfen CR, Zhuang X, Wang Y, Rubin GM, Looger LL (2015) High-performance probes for light and electron microscopy. Nat Methods 12:568–576.

Vizi E, Fekete A, Karoly R, Mike A (2010) Non-synaptic receptors and transporters involved in brain functions and targets of drug treatment. British Journal of Pharmacology 160:785– 809.

Wickham H (2016) ggplot2: Elegant Graphics for Data Analysis. Springer-Verlag New York. Available at: https://ggplot2.tidyverse.org.

Wickham H et al. (2019) Welcome to the Tidyverse. Journal of Open Source Software 4:1686.

Wildenberg G, Sorokina A, Koranda J, Monical A, Heer C, Sheffield M, Zhuang X, McGehee D, Kasthuri B (2021) Partial connectomes of labeled dopaminergic circuits reveal non-synaptic communication and axonal remodeling after exposure to cocaine. eLife 10:e71981.

Williams JL, Shearin HK, Stowers RS (2019) Conditional Synaptic Vesicle Markers for Drosophila. G3 (Bethesda) 9:737–748.

Wu C-L, Shih M-FM, Lee P-T, Chiang A-S (2013) An Octopamine-Mushroom Body Circuit Modulates the Formation of Anesthesia-Resistant Memory in Drosophila. Current Biology 23:2346–2354.

Wu M, Nern A, Williamson WR, Morimoto MM, Reiser MB, Card GM, Rubin GM (2016) Visual projection neurons in the Drosophila lobula link feature detection to distinct behavioral programs Scott K, ed. eLife 5:e21022.

Yan W, Lin H, Yu J, Wiggin TD, Wu L, Meng Z, Liu C, Griffith LC (2023) Subtype-Specific Roles of Ellipsoid Body Ring Neurons in Sleep Regulation in Drosophila. J Neurosci 43:764– 786.

You I-J, Wright SR, Garcia-Garcia AL, Tapper AR, Gardner PD, Koob GF, David Leonardo E, Bohn LM, Wee S (2016) 5-HT1A Autoreceptors in the Dorsal Raphe Nucleus Convey Vulnerability to Compulsive Cocaine Seeking. Neuropsychopharmacol 41:1210–1222.

Young J m., Armstrong J d. (2010) Structure of the adult central complex in Drosophila: Organization of distinct neuronal subsets. Journal of Comparative Neurology 518:1500– 1524.

Yuan Q, Joiner WJ, Sehgal A (2006) A Sleep-Promoting Role for the Drosophila Serotonin Receptor 1A. Current Biology 16:1051–1062.

Zeng J, Li X, Zhang R, Lv M, Wang Y, Tan K, Xia X, Wan J, Jing M, Zhang X, Li Y, Yang Y, Wang L, Chu J, Li Y, Li Y (2023) Local 5-HT signaling bi-directionally regulates the coincidence time window for associative learning. Neuron 111:1118–1135.e5.

